# Loss-of-function mutations in *ASIP* and *MC1R* are associated with coat colour variation in marsupials

**DOI:** 10.1101/2025.01.22.633987

**Authors:** Ryan Sauermann, Bronwyn Fancourt, Tim Faulkner, Hayley Shute, Dean Reid, Andrew J. Pask, Charles Y. Feigin

**Affiliations:** School of BioSciences, The University of Melbourne, VIC, Australia; School of Environmental and Rural Science, University of New England, Armidale, NSW, Australia; Queensland Parks & Wildlife Service, Department of the Environment, Tourism, Science & Innovation, Toowoomba, QLD, Australia; Australian Reptile Park & Aussie Ark, Somersby, NSW, Australia; Department of Sciences, Museums Victoria, Carlton, VIC Australia; Department of Ecological, Plant and Animal Sciences, La Trobe University, Bundoora, VIC, Australia

## Abstract

Pigment production in mammalian hair follicles is governed in part by interactions between Agouti Signalling Protein (ASIP) and the Melanocortin 1 Receptor (MC1R). The most common coat colours in mammals result from alternating bands of dark eumelanin and light pheomelanin within individual hairs. However, coats dominated by a single type of melanin have arisen several times. Here, we examine the genetic basis of two instances among the marsupials: a melanistic morph of the eastern quoll (*Dasyurus viverrinus*) found at high frequency in the wild, and a rare case of fixed xanthism in the marsupial moles. In the eastern quoll, we show that a deletion encompassing part of the *ASIP* coding sequence likely explains melanism in this species. Notably, this mutation is convergent with that recently discovered in its dark-coated relative, the Tasmanian devil (*Sarcophilus harrisii*). Conversely, we show that a nonsense mutation which severely truncates *MC1R* in the southern marsupial mole (*Notoryctes typhlops*) is a strong candidate driver of its pheomelanin-predominant coat. Together with other recent findings, our results suggest that loss-of-function mutations have occurred repeatedly within the marsupials, representing an important mechanism underpinning coat colour variation.

## Introduction

Coat colour in mammals results from the quantities and spatial distributions of two types of pigment, dark eumelanin and light pheomelanin[1]. Within the hair follicle, these pigments are regulated in part by secretion of Agouti Signalling Protein (ASIP) from dermal papilla (DP) cells, which acts in a paracrine manner on pigment-producing melanocytes by binding to the Melanocortin 1 Receptor (MC1R)[2]. MC1R constitutively promotes the production of dark eumelanin, a function that is enhanced by its agonist, α-MSH. ASIP functions as an inverse agonist, both competing with α-MSH to bind MC1R and reducing MC1R’s activity, leading to a shift toward production of light pheomelanin[2]. The most common dorsal coat colours in mammals, ranging from grey to light brown, result from alternating bands of pheomelanin and eumelanin within individual hair shafts, a phenotype called agouti (figure 1a-c)[1]. By contrast, patterns that span many follicles, including stripes and spots, are regulated by mechanisms that establish spatial information across whole skin regions, such as reaction-diffusion systems and long-range morphogens[3].

**Figure 1.**
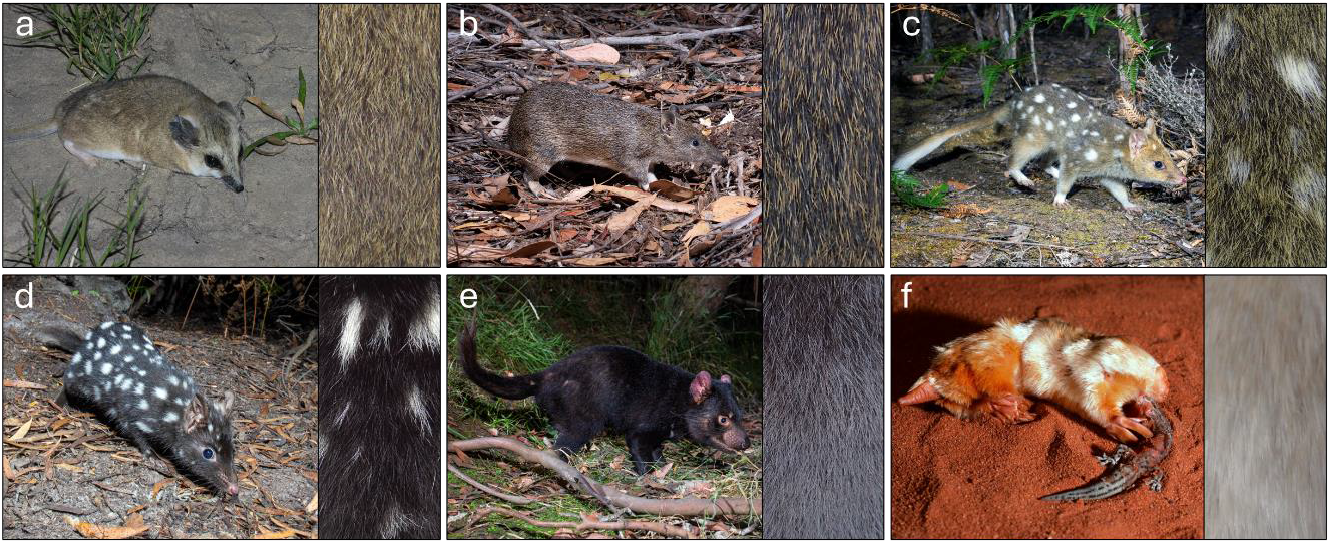
Illustration of marsupial coat colour variation. Species such as the fat-tailed dunnart (*Sminthopsis crassicaudata*; a) and southern brown bandicoot (*Isoodon obesulus*; b) exhibit the typical mammalian agouti pattern with alternating bands of pheomelanin and eumelanin. The eastern quoll (*Dasyurus viverrinus*) has two common colour morphs, a wildtype “fawn” morph (c) with an agouti background coat and a melanistic “black” morph (d). The Tasmanian devil (*Sarcophilus harrisii*; e) a solid black background coat. The marsupial moles (illustrated here by the southern species, *Notoryctes typhlops*; f) typically have a pale-yellow coat. Credits for animal photographs: Lachlan Copeland (a), Brett Vercoe (b-e), Mike Gillam (f).

Loss of a single type of melanin can arise through mutations that modify or inhibit regulators of pigment production. Among the most common examples in mammals is melanism, a phenotype in which eumelanin is produced exclusively or in elevated quantities, leading to dark brown or black hair. Genetic studies have linked several cases of melanism with either loss-of-function (LoF) mutations in *ASIP*, or gain-of-function (GoF) mutations in *MC1R*[4-6]. Conversely, coat phenotypes arising from overproduction of pheomelanin appear to be rarer, or more poorly documented in wild mammals[7, 8]. Such phenotypes can be classified as xanthism or xanthochromism when pheomelanin is yellow in colour, or erythrism when it has a more orange to red hue[9]. Better documented are cases of leucism and albinism, which involve loss of pigmentation. While often considered chromatic disorders[8], these phenotypes are not intrinsically deleterious and their fitness consequences depend on ecological context.

Coats dominated by one type of melanin have arisen multiple times in marsupials, including instances where they have become fixed in a species (figure 1d-f). For example, we recently showed that in the Tasmanian devil (*Sarcophilus harrisii*), a LoF mutation in *ASIP* likely contributes to the species’ distinctive, black coat (figure 1e)[10]. Interestingly, a similar phenotype exists as a high frequency polymorphism in the closely related and parapatric eastern quoll (*Dasyurus viverrinus*; figure 1c,d)[11]. Additionally, the marsupial moles (Genus *Notoryctes*), which comprise two cryptic, fossorial species native to Australia’s great deserts, exhibit a cream to pale yellow colour (figure 1f)[12]. This phenotype represents a rare case of fixed xanthic coloration in a wild mammal.

Here, we examine the genomic basis of coat colour in the eastern quoll and southern marsupial mole. First, using a new haplotype-phased assembly of the eastern quoll and resequencing of phenotyped individuals, we show that a deletion at the *ASIP* locus likely underpins its melanistic morph. Interestingly, this mutation is convergent with that found the closely related Tasmanian devil[10]. Further, by examining the *MC1R* locus across agreodont marsupials (comprising the orders light-yellow, Peramelemorphia and Notoryctemorphia)[13, 14] we identify a nonsense mutation that severely truncates the receptor as a candidate for the marsupial mole’s distinctive light yellow coat. Our results align with several recent findings in other marsupials, illustrating that LoF changes in core pigment regulating genes are potentially important mechanisms for the evolution of coat colour in this lineage.

## Results

### Haplotype-phasing identifies a deletion at the eastern quoll *ASIP* locus

We recently generated a reference genome for the eastern quoll (DasViv_v1.0)[10]. This animal exhibited the wildtype “fawn” colour morph, which is characterized by agouti banded background fur, punctuated by white (pigment-free) spots (figure 1c). A melanistic morph, which exhibits black background fur while retaining white spots (figure 1d), is also found at high frequency in both wild and captive populations, occurring at a ratio of approximately 1:3 relative to the fawn morph[15-17]. This has lead to the hypothesis that melanism in eastern quolls has a recessive pattern of inheritance[17]. Given that *ASIP* LoF is a driver of melanism in other mammals and is expected to be recessive based on its molecular function, we first asked whether our fawn-coloured reference individual may have been a carrier for such a mutation. To this end, we generated a new, haplotype-phased genome of the eastern quoll by reassembling PacBio HiFi and Omni-C reads that were used to produce the previous pseudohaplotype genome. The two resulting haplotypes were named DasViv_v2.0_hap1 and DasViv_v2.0_hap2 (hereafter, referred to as haplotype 1 and haplotype 2, respectively). Scaffold metrics for each haplotype were comparable to those of DasViv1.0 (table S1).

To facilitate examination of the *ASIP* locus, we next annotated this gene in each eastern quoll haplotype assembly, as well as assemblies of several related marsupials (table S2). Interestingly, while the *ASIP* allele annotated on haplotype 1 contained a complete coding sequence as in other marsupials with typical agouti banded hair, annotation of the first two coding exons on haplotype 2, including the start codon, was unsuccessful (data S1). Blastn searches for these exons similarly returned no hits against haplotype 2[18]. Given this, we next extracted the genomic region surrounding the expected locations of these exons from both eastern quoll haplotypes. Alignment of these sequences revealed a ∼4.7 kb stretch of sequence missing in eastern quoll haplotype 2 and encompassing the first two coding exons (figure 2, data S2). This observation indicated that the individual used to generate the eastern quoll reference genome was indeed heterozygous for a presumptive LoF mutation at the *ASIP* locus. This, together with the animal’s fawn colour, is also consistent with melanism having a recessive pattern of inheritance. Interestingly, the eastern quoll *ASIP* deletion appears to be convergent with the fixed *ASIP* deletion recently identified in a closely related dasyurid marsupial, the Tasmanian devil. Both deletions share an extremely similar 5’ break point, though the devil mutation only encompasses the first coding exon (data S2)[10].

**Figure 2.**
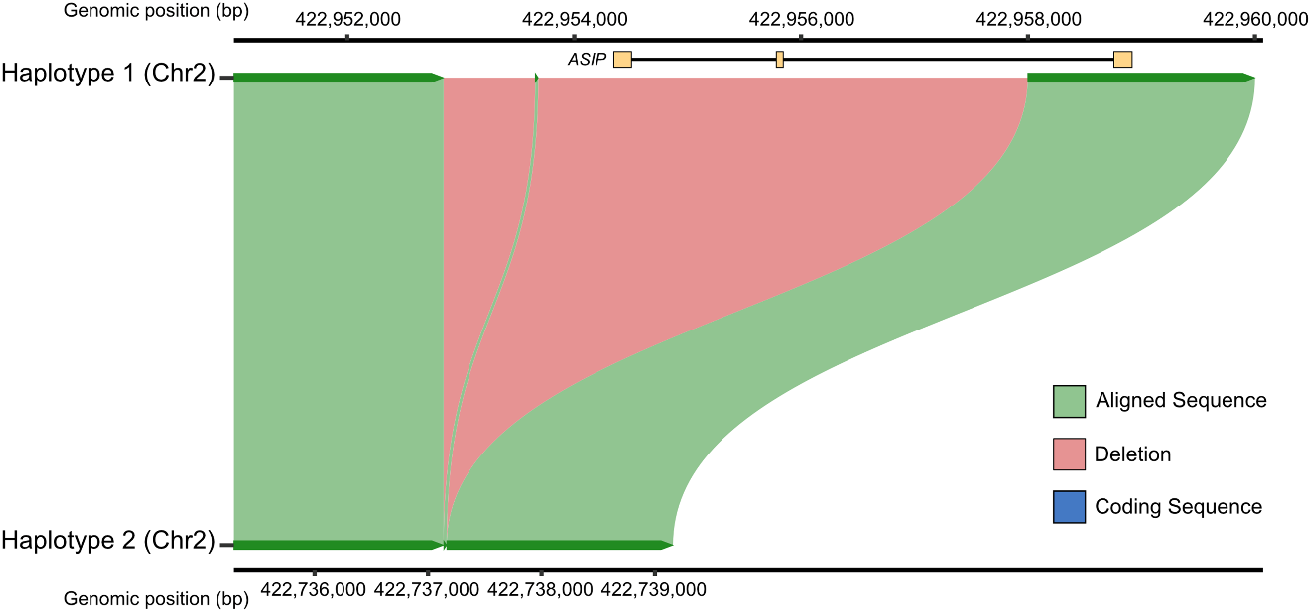
Alignment of the eastern quoll *ASIP* locus, centred on the deletion region. Green ribbons represent sequences aligned between haplotype 1 (top) and haplotype 2 (bottom). Red ribbons represent the putative deletion region, which is present in haplotype 1 but absent from haplotype 2. Coding exons (i.e. CDS annotations) for *ASIP* are shown in yellow above the alignment, illustrating that the first two coding exons are deleted.

### Resequencing supports recessive melanism caused by *ASIP* deletion

To further explore this finding, we sequenced genomic DNA from fawn (n = 3) and black morph (n = 3) eastern quolls and visualized mapping coverage against eastern quoll haplotype 1 (which retains the intact *ASIP* locus). This revealed that all black morph quolls showed zero mapping coverage over the genomic interval that corresponded to the *ASIP* deletion on eastern quoll haplotype 2, while all fawn morph quolls showed high coverage (figure 3). This indicated that all black morph quolls are homozygous for this deletion, reinforcing that a recessive LoF mutation likely explains melanism in eastern quolls.

**Figure 3.**
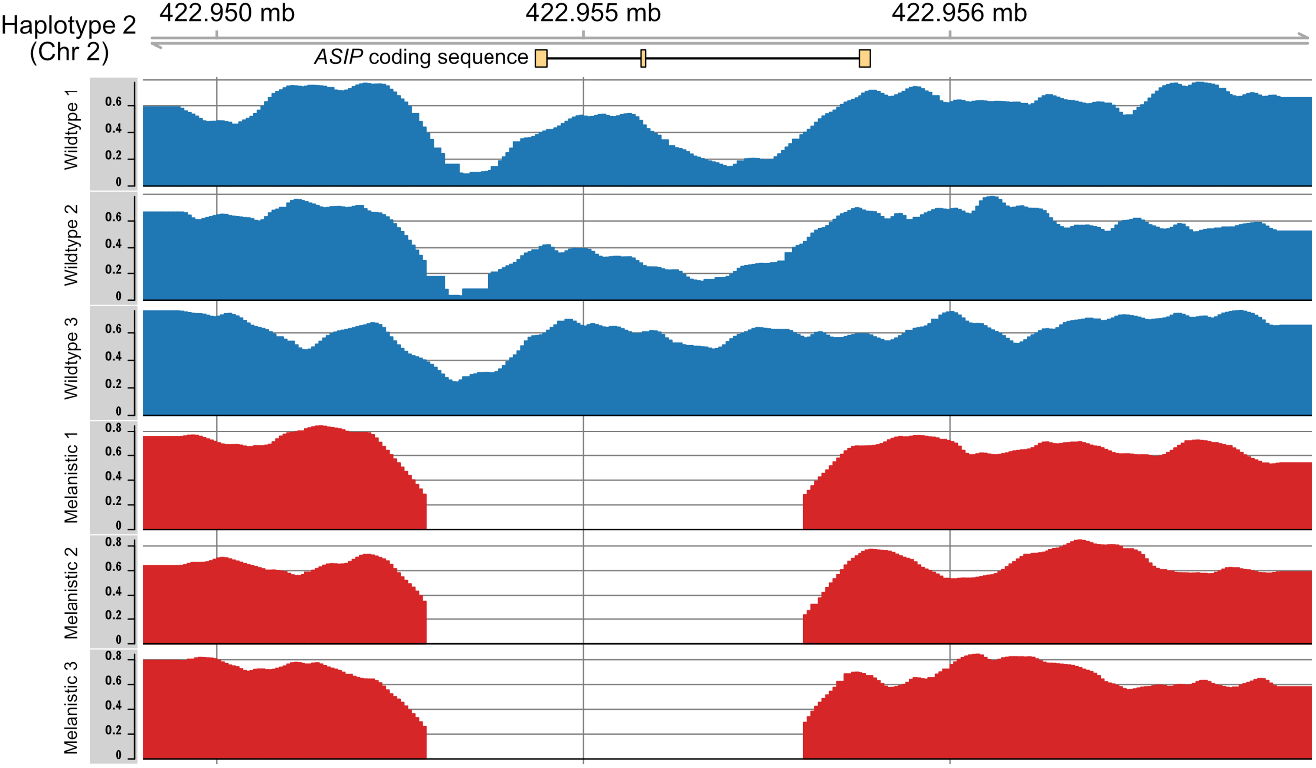
Read mapping coverage of wildtype and melanistic eastern quolls across the *ASIP* locus. Melanistic quolls (red) show no read coverage across the 4.7kb deletion region, indicating homozygosity for this mutation whereas all wildtype quolls exhibit high coverage.

### Truncation of MC1R likely drives yellow coat colour in marsupial moles

Another notable instance among agreodonts where agouti banding has been lost is in the marsupial moles. Comprising two species, the northern and southern marsupial moles (*Notoryctes caurinus* and *Notoryctes typhlops*, respectively), this desert-dwelling group has evolved a suite of convergent adaptations with eutherian moles to facilitate its fossorial lifestyle, including highly reduced eyes and modified forelimbs which they use to “swim” through loose sands[13, 19]. They are among the most rarely observed mammals in Australia, with only a handful of reported sightings each decade. All documented individuals have been found to exhibit a cream to pale yellow pelage, with no obvious sign of dark eumelanin deposition (figure 1f)[20]. We recently generated a reference genome for the southern marsupial mole and thus sought to explore the basis of this species’ unique coat colour[13]. Given the constitutive function of MC1R in eumelanin production, we hypothesized that a LoF at this locus might contribute to its apparent absence in marsupial mole hair.

As *MC1R* was successfully annotated in all marsupial genomes tested, we translated and aligned the amino acid sequences of its orthologs. Interestingly, we found that while *MC1R* was overall highly conserved across these species, a single gap was present, exclusively in the marsupial mole. Examination of the nucleotide alignment at this position showed that the gap was due to an in-frame stop codon at position 160, in place of a highly conserved arginine residue (R160X; figure 4a, data S3). Based on the UniProt structural annotation of mouse MC1R (A0A0B6VTJ4), truncation from the homologous residue (R158 in mouse) is expected to ablate ∼50% of the protein’s length, including 4 of 7 transmembrane domains (figure 4b)[21, 22].

**Figure 4.**
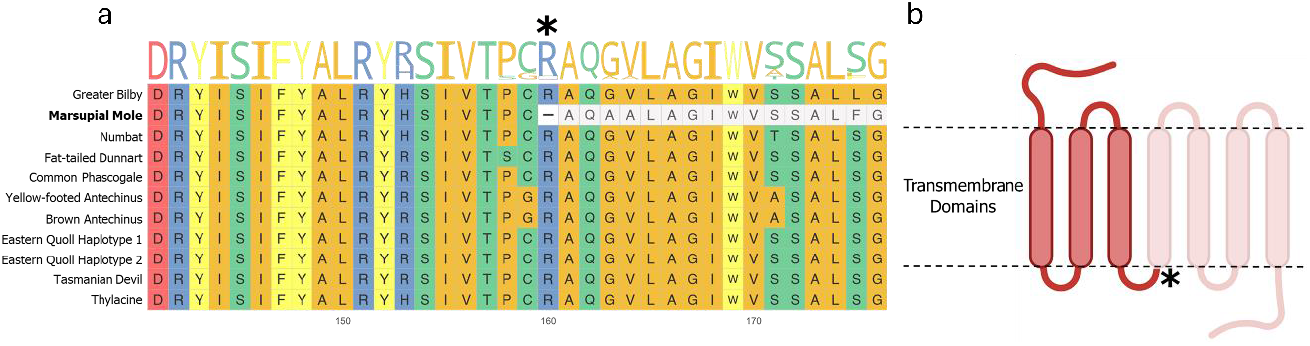
(a) Partial view of aligned amino acid sequences of MC1R orthologs across examined marsupials. Colours indicate residues with similar side chain properties. The sequence logo represents the frequency of different residues at each position. The asterisk indicates the location of the identified nonsense mutation in the marsupial mole, with the truncated bases shown in grey. (b) A diagram of the MC1R protein showing domains ablated by this change.

## Discussion

Here, we provide evidence that repeated loss-of-function mutations in core pigment regulating genes have contributed to coat colour variation within and between marsupial species. We showed that in eastern quolls, a deletion in the *ASIP* locus is associated with the common melanistic morph. Ablation of the only annotated start codon and two complete coding exons likely prevents translation of a functional protein product, resulting in a complete loss of ASIP’s function as an inverse agonist of MC1R. *ASIP* LoF mutations have previously been associated with melanism in various other mammals[4-6]. Intriguingly, this mutation appears to be convergent with a similar deletion recently identified in its close relative, the Tasmanian devil (data S1 & 2)[10]. While differing in size (the devil deletion encompasses only the first coding exon), we expect the effects of these mutations to be similar, due to the loss of the start codon. Of note, the 5’ breakpoints of the Tasmanian devil and eastern quoll deletions are very close in position (data S2). Sequence similarity between genomic regions flanking the deletion region on each haplotype, including several species-specific indels, strongly suggests that the eastern quoll mutation occurred independently from that in the Tasmanian devil. Thus, their deletions are likely convergent, rather than reflecting a shared ancestral variant. As the devil and quoll are closely related and dasyurid genome structure is highly conserved, sequence composition at this locus may have contributed to such similar, convergent deletions occurring in these sister genera[23].

Dark pelage has evolved in other marsupials, including brushtail possums (*Trichosurus vulpecula*), a phenotype recently shown to be driven by a mutation in the *ASIP* coding sequence[24]. The co-occurrence of multiple dark-coated species in Tasmania was noted by Bond et al. 2024, with several possible explanations explored[24]. One hypothesis proposed was a fitness benefit for darker fur colour in Tasmania’s humid climates, in line with Gloger’s rule[25]. Clear data on the frequency of fawn and black morph quolls across Tasmanian environments is not currently available. However, Tasmania as a whole became separated from mainland Australia around ∼14,000 years ago at the end of the Last Glacial Maximum (LGM), and represents the Southern end of the species’ historical range[26]. Black morph eastern quolls were readily observed across Southeastern Australia prior to their extirpation from the mainland in the late 20^th^ century[17, 27]. Presuming the genetic basis of melanism was shared between mainland and modern Tasmanian populations, it is likely that black morph eastern quolls persisted in quite diverse environments and through periods of climatic change. Thus, it remains unclear whether Gloger’s rule provides a sufficient explanation for its relative abundance.

An alternative explanation proposed was crypsis, for example to avoid predation by the recently extinct Tasmanian tiger or thylacine (*Thylacinus cynocephalus*), reflecting an example of the “ghosts of predators past”[24, 28]. A similar hypothesis was recently proposed to explain the balanced frequencies of two colour morphs, green and olive, in the flightless Kākāpō parrot of New Zealand[29]. Unlike the Kākāpō colour polymorphism though, eastern quolls in Tasmania and the historic mainland populations show a roughly 3:1 ratio of fawn and black animals among both adults and pouch young, in line with mendelian expectations[17]. This does not rule out an advantage for dark fur and indeed it is quite notable that an extremely similar deletion at the *ASIP* locus has become fixed in the Tasmanian devil, potentially implying selection in favour of dark pelage. Both species may have been preyed upon by the thylacine, and as Bond et al. 2024 notes, the Tasmanian devil itself may continue to impose similar pressures on eastern quolls. Future population-scale studies of eastern quolls, including historical specimens prior to the extinction of the thylacine and the collapse of Tasmanian populations driven by devil facial tumour disease[30], may provide further insights.

In stark contrast to the eastern quoll and Tasmanian devil, marsupial moles appear to have lost eumelanin production in their fur altogether, a rare phenotype in mammals. Our analyses indicate that an early stop codon that truncates ASIP’s receptor, MC1R, is likely a contributing factor. Experiments using CRISPR-Cas9 to truncate the rabbit ortholog of *MC1R* result in a light yellow coat, due to the loss of eumelanin deposition into the hair shaft[31]. This strongly suggests that the natural truncation of marsupial mole MC1R contributes to its distinctive coat colour. Rabbits in which *MC1R* has been knocked out appear to exhibit a fairly striking yellow hue, reminiscent of spontaneous xanthism/xanthochromism observed in various wild mammals[7-9, 31], whereas the dorsal fur of marsupial moles is a paler yellow or cream colour (figure 1f). This suggests that in addition to the loss of eumelanin production, marsupial moles may also exhibit a degree of hypopigmentation. Pseudogenization of *MC1R* has previously been linked to leucism in cave fish, but not to our knowledge in mammals. Therefore, mutations at other loci involved in pigmentation may contribute to the marsupial mole’s overall coat colour[32]. Specific regions of the marsupial mole’s fur, such as the anterior aspect of the forelimbs and posterior aspect of the hindlimbs, show prominent yellow coloration more typical of xanthism in other species (figure 1f, figure S1). Colour variation across the marsupial mole’s coat may therefore reflect subtle developmental patterning. Notably, the genus *Notoryctes* comprises two distinct species, the southern marsupial mole (examined here) and northern marsupial mole, both of which share a similar coat colour[20]. Molecular studies indicate that these sister species diverged around 4.6 million years ago[19], strongly suggesting that the loss of eumelanin production through MC1R LoF may have arisen prior to their divergence.

Xanthism and leucism are rare in wild mammals, likely due to a fitness cost for conspicuous coloration and are considered chromatic disorders[7, 33]. Thus, for such a phenotype to be fixed in marsupial moles is remarkable. While these animals are fossorial, their lighter coloration may provide some camouflage (depending on the colour of local sands) or may contribute somewhat to thermoregulation when the animal briefly emerges on the surface. While such periods may be short, predation risk or sun exposure may provide some benefit to their coat colour.

Our results suggest that loss-of-function mutations in the pigment regulators *ASIP* and *MC1R* contribute to coat colour in the eastern quoll and marsupial mole respectively. Taken together with recent findings in related marsupials, they suggest that loss-of-function mutations in core pigment regulation genes may represent an important mechanism for the evolution of coat colour in this lineage. Our work further invites new investigations into the potential adaptive implications of such mutations.

## Methods

### Genome Assembly

PacBio HiFi (∼97.68 gigabases) and Omni-C (∼126 gigabases) reads (NCBI BioProject PRJNA75870) previously used to produce a pseudohaplotype assembly of a female, fawn morph eastern quoll were reanalysed to produce a new haplotype-phased assembly[10]. Briefly, PacBio reads were provided as input to hifiasm v0.15.4-r343 with default parameters to produce primary scaffolds[34]. Next, Omni-C reads were aligned against the assembly with bwa mem v0.7.17. The alignment and assembly were then provided to the HiRise pipeline (Dovetail Genomics; Scotts Valley, CA USA) yielding two haplotype assemblies[35]. Assembly quality for each haplotype was compared to that of the original pseudohaplotype genome using BUSCO v5.8.2 (parameters -m genome -l mammalia_odb10) and the stats.sh script provided by the bbmap package with default parameters (table S1)[36, 37]. Scaffold metrics and BUSCO gene recovery were roughly the same between the original pseudohaplotype assembly (DasViv_v1.0) and each haplotype assembly (DasViv_v2.0_hap1 and DasViv_v2.0_hap2, respectively) with marginal improvements in ‘Complete Single Copy’ and ‘Missing’ ortholog categories when the complete haplotype-phased assembly is taken together.

### Identification and alignment of *ASIP* and *MC1R* orthologs

To identify orthologs of *ASIP* and *MC1R* across agreodont marsupials (table S2)[10, 30, 38-43], we used the program LiftOff v1.6.3[44] (default parameters) to perform lift-over annotations. To avoid potential biases introduced by using a reference species with divergent pigmentation, we used the high-quality RefSeq gene models from the agouti-patterned yellow-footed antechinus (*Antechinus flavipes*)[38]. CDS sequences of *ASIP* and *MC1R* were then extracted from each genome using Gffread v0.9.12[45] and aligned using the MAFFT web server (https://mafft.cbrc.jp/alignment/software/)[46]. Searches for *ASIP* coding regions missing in eastern quoll haplotype 2 were performed using Blastn v2.16.0+[18]. As no hits were returned, no results are shown. MC1R orthologs from all examined marsupial genomes were translated and realigned with MAFFT. Visualization of the MC1R amino acid alignment was performed using the R package ggmsa v1.13.1[47].

### Alignment and visualization of the *ASIP* genomic locus

Using the location of annotated *ASIP* exons identified by LiftOff, we extracted the *ASIP* locus from both eastern quoll haplotypes as well as the Tasmanian devil, which were aligned using the MAFFT web server as above. To produce the visualization shown in figure 2, the *ASIP* locus in eastern quoll haplotypes 1 and 2 were realigned with minimap2 v2.28-r1209 (parameters: -x asm20 --eqx --secondary=no -c) and plotted using the R package SvbyEye v0.99.0[48, 49].

### Eastern quoll resequencing

Tissue samples of six eastern quolls of known colour morph (n = 3 fawn, n = 3 black; table S3) were acquired as secondary use from the Aussie Ark captive breeding sanctuary (Somersby, NSW, AU) and through prior field collections on Bruny Island under University of Tasmania Animal Ethics Committee Ethics Approval Permit A11655 and Department of Primary Industries, Parks, Water and Environment permits FA10042 and FA11050. DNA was extracted with the DNeasy Blood & Tissue Kit (Qiagen, Cat. No. 69504). Libraries were prepared using the VAHTS Universal Pro DNA Library Prep Kit for Illumina (Vazyme Cat. No. ND610) and were sequenced to a depth of at least 25X coverage on an Illumina NovaSeq S4 flow cell by Azenta Life Sciences (Burlington, MA, USA).

Raw reads were filtered for quality and residual adaptors using fastp (default parameters) and then mapped against DasViv_v2.0_hap1, which retains a complete copy of the *ASIP* locus, using bwa-mem2 v 2.2.1 (parameter -M)[50]. Alignments were filtered using samtools view v1.13 to retain only primary alignments with a MAPQ greater than 30 and properly paired reads (parameters -f 3 -F 3340 -q 30)[51]. Normalization for mapping coverage and conversion to bedgraph format were performed using deepTools bamCoverage v3.5.5 (−-scaleFactor 10 -- normalizeUsing BPM --exactScaling --normalizeUsing BPM)[52]. Visualization of read coverage over the region shown in figure 3. was performed using the R package Gviz v1.50.0[53].

## Data Availability

Eastern quoll haplotype assemblies 1 and 2 are available on NCBI under BioProjects PRJNA1209419 and PRJNA1209418, respectively. Whole genome resequencing reads are available under BioProject PRJNA1209406. All original code used in this study is attached as supplementary material in code S1-S3. The marsupial mole photograph used in figure 1F is copyrighted (Mike Gillam/AUSCAPE, all rights reserved) and included here on a single use, non-exclusive license.

## Contributions

C.Y.F. conceived the study and wrote the manuscript with editing by all co-authors. C.Y.F. and R.S. performed computational analyses. R.S. processed samples. B.F. provided subsamples of wild individuals. T.F. H.S. and D.R. provided subsamples of captive individuals. A.J.P. provided computational resourcing and sequencing support.

## Acknowledgements

This study was supported by a Wild Genomes grant from Revive & Restore (Contract no. 2020-017) and through philanthropic support by the Wilson Family Trust. We thank Cantata Bio LLC for assistance producing the haplotype-phased eastern quoll genome, Rodrigo Hamede and Christopher Burridge of the University of Tasmania for help with sample logistics, Lachlan Copeland, Brett Vercoe and Mike Gillam for the use of marsupial photographs, Museums Victoria for permission to photograph pelts and Elise Ireland for extensive proofreading.

## Notes

https://github.com/charlesfeigin/ASIP-MC1R-Repository

### Competing Interest Statement

The authors have declared no competing interest.

### Summary of Updates

This version has been updated to fix several minor formatting and spelling errors, to replace the citation of a preprint that was mistakenly included instead of a citation of the published version and to update the preprint's format for a journal submission.

