## Supplementary material for "Loss-of-function mutations in *ASIP* and *MC1R* are associated with coat colour variation in marsupials": tables S1-3, data S1-3, figure S1, code S1-3

**List of Electronic Supplementary Information (ESM)**

Supplementary Tables

table S1: Eastern quoll genome assembly metrics

table S2: Reference genomes of marsupials used in the present study

table S3: Eastern quoll whole genome resequencing

Supplementary Data

data S1: Alignment of agreodont ASIP CDS sequences

data S2: Alignment of the ASIP deletion region

data S3: Alignment of agreodont MC1R amino acid sequences

Supplementary Figures

figure S1: Photos of coat colour in marsupial moles.

Supplementary Code

code S1: Script to plot region alignment between eastern quoll haplotypes at the ASIP locus

code S2: Script to plot eastern quoll mapping coverage across the ASIP locus

code S3: Script to plot amino acid alignment of agreodont MC1R orthologs

| **table S1: Eastern quoll genome assembly metrics** | | | | |
|---|---|---|---|---|
| **Scaffold Metrics** | **DasViv_v1.0** | **DasViv_v2.0_hap1** | **DasViv_v2.0_hap2** | |
| Total Length (bp) | 3023102958 | 3026165475 | 3023198115 | |
| Gap Percentage | 0.001% | 0.003% | 0.003% | |
| G+C Percentage | 36.2% | 36.2% | 36.2% | |
| Chr1 Length (bp) | 714541192 | 715363772 | 715657459 | |
| Chr2 Length (bp) | 662896076 | 662960542 | 663778599 | |
| Chr3 Length (bp) | 628486596 | 627474335 | 626338559 | |
| Chr4 Length (bp) | 468983349 | 468863422 | 468127937 | |
| Chr5 Length (bp) | 292278231 | 293124788 | 292657180 | |
| Chr6 Length (bp) | 255917514 | 258378616 | 256638381 | |
| ChrX Length (bp) | 95405421 | 95675554 | 89374318 | |
| **BUSCO mammalia_odb10 recovery (n: 9226)** | **DasViv_v1.0** | **DasViv_v2.0_hap1** | **DasViv_v2.0_hap2** | **DasViv_v2.0_combined** |
| Complete Single Copy | 8804 (95.4%) | 8774 (95.1%) | 8781 (95.2%) | 8817 (95.6%) |
| Duplicated | 93 (1.0%) | 98 (1.1%) | 92 (1.0%) | 102 (1.1%) |
| Fragmented | 85 (0.9%) | 88 (1.0%) | 88 (1.0%) | 96 (1.0%) |
| Missing | 244 (2.6%) | 266 (2.9%) | 265 (2.9%) | 238 (2.6%) |

| table S2: Reference genomes of marsupials used in the present study | | | |
|---|---|---|---|
| **Species** | **Common Name** | **Assembly** | **Accessed From** |
| *Antechinus stuartii* | Brown antechinus | GCA_016696395.1_USYD_AStu_M | NCBI |
| Phascogale tapoatafa | Brush-tailed phascogale | Phascogale_tapoatafa_HiC | DNAZoo |
| *Dasyurus viverrinus* | Eastern quoll | GCA_020854095.1_UniMelb_DasViv_v1.0 | NCBI |
| *Dasyurus viverrinus* | Eastern quoll | DasViv_2.0_hap1 | Present Study |
| *Dasyurus viverrinus* | Eastern quoll | DasViv_2.0_hap2 | Present Study |
| Sminthopsis crassicaudata | Fat-tailed dunnart | Dunnart_TGS_ONT-PacBio | Ibeh et al. 2024 (*Gigabyte* ) |
| Macrotis lagotis | Greater bilby | GCA_037893015.1_bilby.v1.9.chrom | NCBI |
| *Notoryctes typhlops* | Southern marsupial mole | UniMelb_NotTyph_v1_0 | NCBI |
| *Myrmecobius fasciatus* | Numbat | Myrmecobius_fasciatus_HiC | DNAZoo |
| *Sarcophilus harrisii* | Tasmanian devil | GCF_902635505.1_mSarHar1.11 | NCBI |
| *Thylacinus cynocephalus* | Thylacine | GCA_007646695.3_UniMelb_ThyCyn2.0_hybrid_assembly | NCBI |
| *Antechinus flavipes* | Yellow-footed antechinus | GCF_016432865.1_AdamAnt_v2 | NCBI |

| table S3: Eastern quoll whole genome resequencing | | | |
|---|---|---|---|
| **Sample Name** | **Color Morph** | **Source** | **Sequencing Depth (bases)** |
| Wildtype 1 | Fawn | Bruny Island (Wild) | 111807547500 |
| Wildtype 2 | Fawn | Bruny Island (Wild) | 90566055300 |
| Wildtype 3 | Fawn | Bruny Island (Wild) | 120543666300 |
| Melanistic 1 | Black | Aussie Ark (Captive) | 150382818900 |
| Melanistic 2 | Black | Aussie Ark (Captive) | 165093457200 |
| Melanistic 3 | Black | Aussie Ark (Captive) | 111883012200 |

### data S1: Alignment of agreodont *ASIP* CDS sequences

M_lag – Greater bilby (*Macrotis lagotis*)
N_typ – Marsupial mole (*Notoryctes typhlops*)
T_cyn – Thylacine (*Thylacinus cynocephalus*)
M_fas – Numbat (*Myrmecobius fasciatus*)
S_cra – Fat-tailed dunnart (*Sminthopsis crassicaudata*)
P_tap – Brush-tailed phascogale (*Phascogale tapoatafa*)
A_fla – Yellow-footed antechinus (*Antechinus flavipes*)
A_stu – Brown antechinus (*Antechinus stuartii*)
D_viv_hap1 – Eastern quoll (*Dasyurus viverrinus*) haplotype 1
D_viv_hap2 – Eastern quoll (*Dasyurus viverrinus*) haplotype 2
S_har – Tasmanian devil (*Sarcophilus harrisii*)

Deletion

```
#NEXUS


BEGIN DATA;
 DIMENSIONS NTAX=11 NChar=393;
 FORMAT DATATYPE=DNA INTERLEAVE MISSING=-;
[NAme: M_lag         Len:   393  CheCk:    0]
[NAme: N_typ         Len:   393  CheCk:    0]
[NAme: T_cyn         Len:   393  CheCk:    0]
[NAme: M_fas         Len:   393  CheCk:    0]
[NAme: S_cra         Len:   393  CheCk:    0]
[NAme: P_tap         Len:   393  CheCk:    0]
[NAme: A_fla         Len:   393  CheCk:    0]
[NAme: A_stu         Len:   393  CheCk:    0]
[NAme: D_viv_hap1    Len:   393  CheCk:    0]
[NAme: D_viv_hap2    Len:   393  CheCk:    0]
[NAme: S_har         Len:   393  CheCk:    0]

MATRIX
M_lag            ATGACAACTAAGCATCTGCT CCTTCCCTTCCTTTTGTCCT GCCTGTGGTTCCTGGCTGTC TACTGCCACCTGGCTGAGGA AGAGAAATGGAGTAAGGACA
N_typ            ATGACAGCTAAGCATCTGTT CCTTCCCTTCCTTCTGACCT GCCTGTGGTTCCTGACTGTC TATTGCCACCTGGCTGATGA AGAGAAATGGAGTAAGGACA
T_cyn            ATGACAGCTAAGCATCTGCT CCTTCCCTTCCTTCTGGTCT GCCTGTGGTTCCTGGCTGCC TACTGCCACCTGGCTGAGGA AGAGAAATGGAGTAAAGATA
M_fas            ATGACAGCTAAGCATCTGTT CCTTCCCTTCCTTCTGGCCT GCCTGTGGTTCCTGGCTGCC TACTGCCACCTGGCTGAGGA AGAGAAATGGAGTAAGGATA
S_cra            ATGACAGCTAAGCATCTGTT CCTTCCTTTCCTTCTGGCCT GCCTGTGGTTCCTGGCTGCC TACTGCCACCTGGCTGAAGA AGAGAAATGGAGTAAGGATA
P_tap            ATGACAGCTAAGCATCTGTT CCTTCCCTTCCTTCTGGCCT GCCTGTGGTTCCTGGCTGCC TACTGCCACCTGGCTGAGGA AGAGAAATGGAGTAAGGATA
A_fla            ATGACAGCTAAGCATCTGTT CCTTCCCTTCCTTCTGGCCT GCCTGTGGTTCCTGGCTGCC TACTGCCACCTGGCTGAGGA AGAGAAATGGAGTAAGGATA
A_stu            ATGACAGCTAAGCATCTGTT CCTTCCCTTCCTTCTGGCCT GCCTGTGGTTCCTGGCTGCC TACTGCCACCTGGCTGAGGA AGAGAAATGGAGTAAGGATA
D_viv_hap1       ATGACAGCTAAGCATCTGTT CCTTCCCTTCCTTCTGGCCT GCCTGTGGTTCCTGGCTGCC TACTGCCACCTGGCTGAGGA AGAGAAATGGAGTAAGGATA
D_viv_hap2       -------------------- -------------------- -------------------- -------------------- --------------------
S_har            -------------------- -------------------- -------------------- -------------------- --------------------

M_lag            GAGGTCTGGCCAGAAACTCT ACCATGAACCTGCCTGCCTT TCCTTCAGTCTCCATTGTAG CACTGAACAAAAAATCCAAG AAGATCATCAGGAAAGAGAT
N_typ            GGGGTTTGGGGAGAAACTCT ACCATGAACCTGCCTGACTT TCCTTCAGTCTCCATCGTAG CACTGAATAAGAAATCCAAG AAGATCACCAGGAAAGAGAT
T_cyn            GGAGTCTGGGAAGAAACT-- -CCATGAACTTGCCTGATTT TCCTTCTGTATCCATCGTGG CACTGAATAAAAAATCCAAG AAGAGCATCAGGAAAGAGAT
M_fas            AGGGTCTGGGAAGAAATT-- -CCATGAACCTGACTGACTT TCCTTCTGTCTCCATCGTGG CATTGAACAAAAAATCCAAG AAGAGCATCAGGAAAGAGAT
S_cra            GGGTTCTGGGAAGAAGCT-- -CCATGAACCTGCCTGACTT TCCTTCTGTGTCCATTGTGG CATTGAATAAAAAAGCCAAG AAGAACATCAGGAAAGAGAT
P_tap            GGGATCTGGGAAGAAGCT-- -CCATGAACCTGCCTGACTT TCCTTCTGTGTCCATCGTGG CATTGAACAAAAAATCCAAG AAGAGCATCAGGAAAGAGAT
A_fla            GGAGTCTGGGAAGAAGCT-- -CCATGAACCTGCCTGACTT TCCTTCTGTGTCCATCGTGG CACTGAACAAAAAATCCAAG AAGAGCATCAGGAAAGAGAT
A_stu            GGGGTCTGGGAAGAAGCT-- -CCATGAACCTGCCTGACTT TCCTTCTGTGTCCATCGTGG CATTGAACAAAAAATCCAAG AAGAGCATCAGGAAAGAGAT
D_viv_hap1       GGGGTCTGGGAAGAAGCT-- -CCATGAACCTGCCTGACTT TCCTTCTGTGTCCATCGTGG CACTGAACAAAAAATCCAAG AAGAACATCAGGAAAGAGAT
D_viv_hap2       -------------------- -------------------- -------------------- -------------------- --------------------
S_har            -------------------- -------------------- -------------------- CACTGAACAAAAAATCCAAG AAGAACATCAGGAAAGAGAT

M_lag            AGAAACCAAGAAATCCTCTG AGAAAAAGGCCCTGATGAAG AAGAACTCTCGACCCCCACC TCCAGCCAACTGTGTAGCCA CCTGGGCCAACTGCCAGCCC
N_typ            AGAAACCAAGAAATCCTCTG AGAAAAAGGCCCTGGTGAAG AAGAGCCCTCAGCCCCCACC TTCAGCCAACTGTGTGGCCA CCTGGGCCAACTGCCAGCCC
T_cyn            AGAAACCAAGAAATCCTCAG AGAAAAAGGCTGTGGTGAAG AAG---------TCCTCATC TGGATCCAACTGTGCAGCCA CAGGGGCCTTCTGCCAGCCC
M_fas            AGAAACCAAGAAATCCTCCG AGAAAAAGGCCGTGGTGAAG AAG---------TCCCCGTC TAGATCCAACTGCGCGGCCA CGGGGGCCTTCTGCCAGCCC
S_cra            AGAAATCAAGAAATCCTTCG AGAAAAAAGCTGTGGTGAAG AAG---------GCCTCATC TGCATCCAATTGTGCGGCCA CAGGGGCCTTCTGCCAGCCC
P_tap            AGAAACCAAGAAATCCTCCG AGAAAAAAGCTGTGGTGAAG AAG---------TCCTCATC TGCATCCACCTGTGCAGCCA CAGGGGCCTTCTGCCAGCCC
A_fla            AGAAACCAAGAAATCCTCCG AGAAAAAAGCTGTGGTGAAG AAG---------TCCTCATC TGCATCCACCTGTGCAGCCA CAGGGGCCTTCTGCCAACCT
A_stu            AGAAACCAAGAAATCCTCCG AGAAAAAAGCTGTGGTGAAG AAG---------TCCTCATC TGCATCCACCTGTGCAGCCA CAGGGGCCTTCTGCCAGCCT
D_viv_hap1       AGAAACCAAGAAATCCTCCG AGAAAAAAGCTGTGGTGAAG AAG---------TCCTCATC TGCATCCACCTGTGCAGCCA CAGGGGCCTTCTGCCAGCCT
D_viv_hap2       -------------------- --AAAAAAGCTGTGGTGAAG AAG---------TCCTCATC TGCATCCACCTGTGCAGCCA CAGGGGCCTTCTGCCAGCCT
S_har            AGAAACCAAGAAATCTTCCG AGAAAAAACCTGTGGTGAAG ---------------TTATC TGCATCCACCTGTGCAGCCA CAGGGGCCTTCTGCCAGCCT

M_lag            CTGGCATCACCCTGCTGCAA CCCATGTGCCATATGTCACT GCCGCTTCTTCCGTAGTGTC TGCTCCTGCCGCCTGTTCCG GCCCAGTTGCTAA
N_typ            CTGACGTCACCCTGCTGCAA TCCATGTGCTATATGCCACT GCCGCTTCTTCCGCAGTGTC TGCTCCTGCTGCTTGCTCCA GCACAGCTGCTGA
T_cyn            CAGACACTGTCCTGCTGCAA CCGGTGCGCCACATGCCACT GCCGCTTCTTCAGGAGCTCC TGCTCCTGCCGTCTGTTCCA GCCGGGCTGCTGA
M_fas            CACACGCTGTCCTGCTGCGA GCCGTGCGCCTCGTGCTACT GCCGCTTCTTCGGGAGAGTC TGCTCCTGCCGCCTGTTCCA GCAGAGCTGCTGA
S_cra            CAGACAATTTCCTGCTGCAA CCCGTGTGACTCGTGCCAAT GCAGGTTCTTCAGGAGTGCC TGTTCCTGCCGCTCATTCAT GGGGAAATGCTGA
P_tap            CAGACCATTTCCTGCTGCAA CAAGTGTGACACGTGCCACT GCCGCTTCTTCGGGAGCGTC TGCTTCTGCCGCCCATTCAT GCGGAAATGCTGA
A_fla            CAGACCATTTCCTGCTGCGA CAAGTGTGACACGTGCCACT GCCGCTTCTTCGGGAGCGTC TGCTTCTGCCGCCCGTTCAT GCGGAAATGCTGA
A_stu            CAGACCATTTCCTGCTGCGA CAAGTGTGACACGTGCCACT GCCGCTTCTTCGGGAGCGTC TGCTTCTGCCGCCCGTTCAT GCGGAAATGCTGA
D_viv_hap1       CAGACCATTTCCTGCTGCAA CAAGTGTGATACGTGCCACT GTCGCTTCTTCGGAAGTGTC TGCTTCTGCCGCCAGTTCCT GCGGAAATGCTGA
D_viv_hap2       CAGACCATTTCCTGCTGCAA CAAGTGTGATACGTGCCACT GTCGCTTCTTCGGAAGTGTC TGCTTCTGCCGCCAGTTCCT GCGGAAATGCTGA
S_har            CAGACCATTTCCTGCTGCAA CAAGTGTGATACGTGCCACT GTCGCTTCTTCGGAAGTGTC TGCTTCTGCCGCCCGTTCCT GCGGAAATGCTGA

;
END;
```

### data S2: Alignment of the *ASIP* deletion region

D_viv_hap1 – Eastern quoll (*Dasyurus viverrinus*) haplotype 1
D_viv_hap2 – Eastern quoll (*Dasyurus viverrinus*) haplotype 2
S_har – Tasmanian devil (*Sarcophilus harrisii*)

Deletion
Deleted Coding Sequence
Intact Coding Sequence

```
#NEXUS


BEGIN DATA;
 DIMENSIONS NTAX=3 NCHAR=6685;
 FORMAT DATATYPE=DNA INTERLEAVE MISSING=-;
[Name: S_har          Len:  6685  Check:    0]
[Name: D_viv_hap2          Len:  6685  Check:    0]
[Name: D_viv_hap1          Len:  6685  Check:    0]

MATRIX
S_har                 TCTTACCCATCTTTTAATGG GTATTAAATACCGAATTCTC TAG-AAGCCTTCTTTGAAAC AGTCTCCCTTTACACTGTCC ATATCTTGTATTTTCATAAT
D_viv_hap2            TCTTACCCATCTTTTAATGG GTATTAAATACCGAATTCTC TAGAAAGCCTTCTTTGAAAC ATTCTCCCTTTACACTGTCC CTATCTTGTATTTTCATAAT
D_viv_hap1            TCTTACCCATCTTTTAATGG GTATTAAATACCGAATTCTC TAGAAAGCCTTCTTTGAAAC ATTCTCCCTTTACACTGTCC CTATCTTGTATTTTCATAAT

S_har                 TATCTGCAAGTTGCCTCCTC CATTAGAAGGTGAATTCCTT GAGAGTAGGAAATTAGGCTT TTTCCTTTCCTTAAAACCCC ACGGTTAAGGGCAGTGGCTG
D_viv_hap2            TATCTACAAGTTGCCTCCTC CATTAGAAGGTGAATTCCTT GAGAGTAGGAAA-TAGGCTT TTTCCTTTCCTTAAAACCCC AGGGTTAAGGGCAGTGTGTG
D_viv_hap1            TATCTACAAGTTGCCTCCTC CATTAGAAGGTGAATTCCTT GAGAGTAGGAAA-TAGGCTT TTTCCTTTCCTTAAAACCCC AGGGTTAAGGGCAGTGTGTG

S_har                 -------------------- -------------------- -----GCACATAGTATCAAT CAATCAAT------------ --------------------
D_viv_hap2            TGTGTATATATATATATATT TTTACAAATATATA------ -------------------- -------------------- --------------------
D_viv_hap1            TGTGTATATATATATATATA TATATATATATACATACACA CACACACACATAGTATCAAT CAATCAATAAACATTTATTG TGTCTACACACCTGTCAGAT

S_har                 -------------------- -------------------- -------------------- -------------------- --------------------
D_viv_hap2            -------------------- -------------------- -------------------- -------------------- --------------------
D_viv_hap1            TGGCTAAGATGACAGGAAAA AATAATAATGATTGTTGGAG GGGATGTGGGAAAACTGGGA CATTGTTGCATTGTTGGTGG AGTTGTGAACGAATCCAACC

S_har                 -------------------- -------------------- -------------------- -------------------- --------------------
D_viv_hap2            -------------------- -------------------- -------------------- -------------------- --------------------
D_viv_hap1            ATTTTGGAGAGTTGTTTGGA ACTATGCTCAAAAAGTTATC AAACTGTGCATACCCTTTGA TCCAGCAGTGTTACTACTGG GCTTATATCCCAAAGAGATT

S_har                 -------------------- -------------------- -------------------- -------------------- --------------------
D_viv_hap2            -------------------- -------------------- -------------------- -------------------- --------------------
D_viv_hap1            ATAAAGCAGGGAAAGGGACC TGTATGTGCACGAATGTTTG TGGCAGCCCTTTTTGTAGTG GCTAGAAACTGGAAGCTGAA TGGATGCCCATCAGTTGGAG

S_har                 -------------------- -------------------- -------------------- -------------------- --------------------
D_viv_hap2            -------------------- -------------------- -------------------- -------------------- --------------------
D_viv_hap1            AATGGCTGAATAAATTGTGG TATATGAATACTATGGAATA TTACTGTTCTGTAAGAAATG ACCAACAGGATGATTTCAGA AAGGCCTGGAGAGACTTACA

S_har                 -------------------- -------------------- -------------------- -------------------- --------------------
D_viv_hap2            -------------------- -------------------- -------------------- -------------------- --------------------
D_viv_hap1            TGAACTGATGCTGAGTGAAA TGAGCAGGACCAGGAGAACA TTATATACTTCAACAACAAT ACTATATGATGCCCAGTTCT GATGGACCTGGCCATCCTCA

S_har                 -------------------- -------------------- -------------------- -------------------- --------------------
D_viv_hap2            -------------------- -------------------- -------------------- -------------------- --------------------
D_viv_hap1            GCAACGAGATCAACCAAATC ATTTCCAATGGAGCAGTAAT GAACTGAACCAGCTATGCCT AGAGAAAGAACTTTGGGAGA TGACGAAAAACCAATACATT

S_har                 -------------------- -------------------- -------------------- -------------------- --------------------
D_viv_hap2            -------------------- -------------------- -------------------- -------------------- --------------------
D_viv_hap1            GAATTCCCAATCCCTATATT TATGCCCACCTGCATATTTG ATTTCCTCCACAAGCTAATT GCACAATATTTCAGAATCAG ATTCTTTTTATACAGCAAAA

S_har                 -------------------- -------------------- -------------------- -------------------- --------------------
D_viv_hap2            -------------------- -------------------- -------------------- -------------------- --------------------
D_viv_hap1            TATGTTTTGGTCATGAATAC TTATTGTATATCTAATTTAT ATTTTAATGTATTTAACATC TACTGGTCATCCTGCCATCT AGGGGAAGGGGTGGGGGGTG

S_har                 -------------------- -------------------- -------------------- -------------------- --------------------
D_viv_hap2            -------------------- -------------------- -------------------- -------------------- --------------------
D_viv_hap1            GGAGGCGAAAAATTGGAACA AGAAAGTTGGCAATTGTTAA TGCTGTAAAGTTATCCATGC ATATAACCTGTAAATAAAAG GCTATTATTAAAAAAAAATT

S_har                 -------------------- ---AAACATTTATTGTGTCT ATTAAAGTAAGCAATTCATC AATGCTTAATGCCCAGTGAC TAACTGCCCCTCCTCACACC
D_viv_hap2            -------------------- -------------------- -------------------- -------------------- --------------------
D_viv_hap1            TTTTTTTAAATTAAAAAAAA AAAAAACATTTATTGTGTCT ATTAAAGCAAGCAATTCATC AATGCTTAATGCCCCGTGAC TAACTGCCCCTCCTCACACC

S_har                 TTCCA--------------- -------------------- -------------------- -------------------- --------------------
D_viv_hap2            -------------------- -------------------- -------------------- -------------------- --------------------
D_viv_hap1            TTCCACAGTGGTTTATTTAT TTTGGTTCTTTTTAATGCAC ATATTGTGTGATAGTTGTGG GTACCACATCTCCCTTCTAG AACATGTGCTCCCTACAGAT

S_har                 -------------------- -------------------- -------------------- -------------------- --------------------
D_viv_hap2            -------------------- -------------------- -------------------- -------------------- --------------------
D_viv_hap1            GGGATTCAGGAGGTATTTAA TAAATGTTCATTGTATTTGT TGTTCTGACATAGCCAGACC GTAACAGTAAATGCTGCTAA GCAGAGGCCATCCTTCTGTT

S_har                 -------------------- -------------------- -------------------- -------------------- --------------------
D_viv_hap2            -------------------- -------------------- -------------------- -------------------- --------------------
D_viv_hap1            TTCCTCCAGTCCTCCCAGTG TCTTCTCATCTCCCCACAGT TTTTGTCTTTTTCCTTTCCT CCCACTGCACTTGTCTCCCC TCCCCTGCTTTAGGGCTAGA

S_har                 -------------------- -------------------- -------------------- -------------------- --------------------
D_viv_hap2            -------------------- -------------------- -------------------- -------------------- --------------------
D_viv_hap1            CCACTCTGTGGCTCATGGGG TCACCATCCAGGGCAGATCT CCCCCACTCCCCCTTCTCCC CAGCCTTCACACATCCCAAT CCCATCTTGACTTCTGGTTT

S_har                 -------------------- -------------------- -------------------- -------------------- --------------------
D_viv_hap2            -------------------- -------------------- -------------------- -------------------- --------------------
D_viv_hap1            TCTGTTCTGCTTAGAAGTTC CAGGATGACAGCTAAGCATC TGTTCCTTCCCTTCCTTCTG GCCTGCCTGTGGTTCCTGGC TGCCTACTGCCACCTGGCTG

S_har                 -------------------- -------------------- -------------------- -------------------- --------------------
D_viv_hap2            -------------------- -------------------- -------------------- -------------------- --------------------
```

```
D_viv_hap1         AGGAAGAGAAATGGAGTAAG GATAGGGGTCTGGGAAGAAG CTCCATGAACCTGCCTGACT TTCCTTCTGTGTCCATCGTG GGTGAGTAGCCTGGCCAGCC

S_har              -------------------- -------------------- -------------------- -------------------- --------------------
D_viv_hap2         -------------------- -------------------- -------------------- -------------------- --------------------
D_viv_hap1         ACTATCTCTGACACAGGACT AGTGTGCAGGGCGGGCCTAT GATCTCTATCCTGGGCTTGG GCTTCGTTTTCCCCTTTTGG GAGATTCCGGGTGGCCTCCC

S_har              -------------------- -------------------- -------------------- -------------------- --------------------
D_viv_hap2         -------------------- -------------------- -------------------- -------------------- --------------------
D_viv_hap1         TCAAATGCAGCTACCTCTCT CAAGTTCTAGCTATTCTTGA TAGTGGTGCTCACTCCCCAT TTAGTCTATAGATTGGGAAC CACAAGAACTGGGTTCCAAT

S_har              -------------------- -------------------- -------------------- -------------------- --------------------
D_viv_hap2         -------------------- -------------------- -------------------- -------------------- --------------------
D_viv_hap1         ACCTGAGAAAATCACTTCTG AAACCCACTCTCAGCATTTC AGATTTCTTCTTATTAAAAA TGGGGAGGGGGGATTTAAAT GCTCTCTAATGTCCTTTCCA

S_har              -------------------- -------------------- -------------------- -------------------- --------------------
D_viv_hap2         -------------------- -------------------- -------------------- -------------------- --------------------
D_viv_hap1         GATCTAAAAATTTTATGATC CCTTCTACTAAATGGCTACC TCCCCTGCCTTTTCTTCAGC TGCTTGCCAGCCTGAAGTCC CTATGGAGTCACAGAGCCTT

S_har              -------------------- -------------------- -------------------- -------------------- --------------------
D_viv_hap2         -------------------- -------------------- -------------------- -------------------- --------------------
D_viv_hap1         ATTAATCAAATCTGGAGAAA AGTCAGATGAATGTAACTCC TTTCTATAGCTAATGTTCAG CATATGAGAGGTGAGCTTTA ATGAGACCTTTGATTCTACC

S_har              -------------------- -------------------- -------------------- -------------------- --------------------
D_viv_hap2         -------------------- -------------------- -------------------- -------------------- --------------------
D_viv_hap1         TGAAAAACAGGGTCTAGACC AATGAGATTTCAGTGCATCT AATATAGATTAAATGTGAAA TTCTTCTAAACATATTTCCA CATTTATCATTCTGCATAAG

S_har              -------------------- -------------------- -------------------- -------------------- --------------------
D_viv_hap2         -------------------- -------------------- -------------------- -------------------- --------------------
D_viv_hap1         AAAAATCAGATCAAAAAGGG GGGGAAATGAGAAAGAAAAA AGCAAGCAAGCAACAAAAAT GAGGGAGAAGCGAGATGGAA AGCAATTTTATTTTGATACA

S_har              -------------------- -------------------- -------------------- -------------------- --------------------
D_viv_hap2         -------------------- -------------------- -------------------- -------------------- --------------------
D_viv_hap1         ATTGACTTCCTTTGCAAATC TTTTTTTTTTCATGAGTTGA AAAACATTGTGCTGGAAAGG GTTTACAGGCTTCGGCCCAG ACTCCAAAAGGGCCCCATGG

S_har              -------------------- -------------------- -------------------- -------------------- --------------------
D_viv_hap2         -------------------- -------------------- -------------------- -------------------- --------------------
D_viv_hap1         CAAAAAAAAAGTTAAGAGCC TATCTAGATAGTGCTGGAGA CTGAGGGATTAGGAAAGGTA GTCTCTAAGACTAGTTTAAA GGTTTCCACAAACCTCTTGG

S_har              -------------------- -------------------- -------------------- -------------------- --------------------
D_viv_hap2         -------------------- -------------------- -------------------- -------------------- --------------------
D_viv_hap1         TCTCAACTGTTCCCAGAACC CAAGGAAAAAAAGGGGGCAA TTGAAACTCCTTTTGACCCT GAGATACTAGGAGGGGAATG CCAAAAGTGGACAAGAATCA

S_har              -------------------- -------------------- -------------------- -------------------- --------------------
D_viv_hap2         -------------------- -------------------- -------------------- -------------------- --------------------
D_viv_hap1         AATTTCTTTTCTGTCTAGCC TGTAAGACTTCCTCTATTCT CAGGCTTTCTCCTCTTTGTT TGTGAGGCACTTACCTCAAT CCCTGGCCTTTGCTCTTTTG

S_har              -------------------- -------------------- -------------------- -------------------- -----------------CAG
D_viv_hap2         -------------------- -------------------- -------------------- -------------------- --------------------
D_viv_hap1         GCTCCTCTAGCATTGAAATG GTCAGGCAAAAATGGTCAGG GGTCTCCACTCCTGCCTCCG GGAGACCCCCTTCCCAGAGG TCTCCCTCTTCCCAATTCAG

S_har              TCTATCCCTGGCCCTGCCAG TCAGACATATCTTCTCTTCA TTTTGTTTTCTTTAAAGCAC TGAACAAAAAATCCAAGAAG AACATCAGGAAAGAGATAGA
D_viv_hap2         -------------------- -------------------- -------------------- -------------------- --------------------
D_viv_hap1         TCTATCCCTGGCCCTGCCAG TCAGACATATCTTCTTTTCA TTTTGTTTTCTTTGAAGCAC TGAACAAAAAATCCAAGAAG AACATCAGGAAAGAGATAGA

S_har              AACCAAGAAATCTTCCGAGG TAGGTGGGACTTCAGCAGAG GTGAAAGAGGGATGGGAGGG GGAGAGGTAAGAGTCCACCC CTGCTTCAAAATAACATGAT
D_viv_hap2         -------------------- -------------------- -------------------- -------------------- --------------------
D_viv_hap1         AACCAAGAAATCCTCCGAGG TAGGTGGGACTTCAACAGAG GTGAAAGAGGGATGGGAGGG GGAGAGGTAAGAGTCCACCC CTGCTTCAAAATAACATGAT

S_har              GGACTGACGTGTGTGTGTGT GTGTTTGTGTGTGTATATAT ATTATAACATGATGAACTGA CT--TGTGTGTGTATAGCAT GATGGATTGA----------
D_viv_hap2         -------------------- -------------------- -------------------- -------------------- --------------------
D_viv_hap1         GGACTGGTGTGTGTGTGTGT G----TGTGTGTGTGTGTGT ATTATAACATGATGAACTGA CTTGTGTGTGTGTATAACAT GATGGATTGATGTGTGTGTG

S_har              -------------------- -------------------- ----------TGTATGTGTG TATATGTATATAGTCTGTAC CCCCCCCCCCCATACTGTCT
D_viv_hap2         -------------------- -------------------- -------------------- -------------------- --------------------
D_viv_hap1         TGTGTATATATATATGTGTG TGTGTGTGTGTGTGTGTGTG TGTGTGTGTGTGTGTGTGTG TGTATGTATATAGTCTGTA- CCCTCCCCCCCATACTGTCT

S_har              GGTGAAGTATGGGGTGGGGA AGGTTGGGTGGGAATATTAA TACCAACATTTTTTCAGGTG AAAACAACTGAGACTTGGAA AGATTGACTTGCTTACAATT
D_viv_hap2         -------------------- -------------------- -------------------- -------------------- --------------------
D_viv_hap1         GGTGAAGTATGGGGTGGGGA AGGTTGGATGGGAATATTAA TACCAACATTTTTTCAGGTG AAAACAACTGAGACTTGGAA AGACTGATTTGCTTACAATT

S_har              ATAATGCCATTAAACAAGAA TCCTATCCAATTTTTATGAC TTGCTCTTTCCAGAATTCTT TCTGTGGTAGCAGTTCTCCA CTCCCTCAAGAAACATATCA
D_viv_hap2         -------------------- -------------------- -------------------- -------------------- --------------------
D_viv_hap1         ATAATGCCATTAAGCGAGAC TCCTATCCAATTTTTATGAC TTGCTCTGTCCAGAATTCTT TCTGTGGTAACAGTTCTCTA CTCCCTCAGGAAACATATCA

S_har              TTGCTTTATTGGTGGAAAGG ATATTAGGAATTGAGAGAAA TGGTTATCTTATATTAATAA CCAACTACTGAATTAGGACT AAGGTTTTTTTCCCTTTTCT
D_viv_hap2         -------------------- -------------------- -------------------- -------------------- --------------------
D_viv_hap1         TTGCTTTATTGGTGGAAAGG ATATTAGGAATTGAGAGAAA TGATTATCTTATATTAATAA CCAACTACTGAATTAGGACT AAGG-TTTTTTCCCTTTTCT

S_har              ACTTCCCCATTCAGAAATCA GCCCTCTTTGGGTTTTTCAG GCAGCCTCTGAAGAAAAGGG CTGATAATATGAGAGAAAAC TCAGATCTCTTCAAAGGATA
D_viv_hap2         -------------------- -------------------- -------------------- -------------------- --------------------
D_viv_hap1         ACTTCCCCATTCAGAAATCA GCCCTCTTTGGGTTTTTCAG GCAGCCTTTGAAGCAAAGGG CTGATAATATGAGAGAAAAC TCAGATCTGTTCAAAGGATA

S_har              AAAGTCATTATAGTTTAAAC AAATTGCCCATTGTATTTTA CTCAAACTTTCCATCCAGGG AAAAAGTGAATACTCCTTTT TATCTAGATTACTCTAGACA
D_viv_hap2         -------------------- -------------------- -------------------- -------------------- --------------------
D_viv_hap1         AAAGTCATTATAGTTTAAAC AAATTGCCCCTTGTATTTTA CTAAAACTTTCCATCCA-GG AAAAAGTGAGTACTCCTTTT TATCTAGACTACTCTAGACA

S_har              GGGTAACATGAACATAGATA AATCTCTTTCACAAAGACTA AAGGATCAGAATCATGTCTC CCTACACCCTCATTTGAATA AGTCCACCTCTTATAATATT
D_viv_hap2         -------------------- -------------------- -------------------- -------------------- --------TTTTACAATATT
D_viv_hap1         GGGTAACATGAACATAGATA AATCTCTTTGACAAAGACTA AAGGATCAGAATCATGTCTC CCTACACCCTCATTTGAATA TGTCCACCTCTTATAATATT

S_har              CATTCTCTCATTACCTG--- ----TTAACCAATCAGATTG GTTAACTCTCCCAAGGGCAT TGAATGGATTCTACAAGATG TCTTTGGCATTGAGAGTGCC
D_viv_hap2         T------------------- -------------------- -------------------- -------------------- --------------------
D_viv_hap1         CATGCTCTCATTACCTGATT GGTAATGGTTAACCAGATTG GTTAACTCTCCCAAGGGCAT TGAATGGATTCTACAAGATG TCTTTGGCATTGAGAGTGCC

S_har              ATTGACCTCTTTTATTAATA ACTGCTAGTATGATTAATAA AATGATTAATTATCCAGAAA CTATTTCTCTTGAACTTTTT ATATGTAACAGATTATTATG
D_viv_hap2         -------------------- -------------------- -------------------- -------------------- --------------------
D_viv_hap1         ATTGACCTCTTTTATTAATT ACTGCTAGTGTGATTAATAA AATGATTAATTACCCAGAAA CTATTTCTCTTGAACTTTTT ATATGTAACAG--TATTTTG

S_har              GAAATATCTTGATTCACTAA AAAAAATATTGCTGCCAATC ATACCTATCAATATATACAG -----AACAGCAAAACCAAG GTTTATTATATGTTTTAACA
D_viv_hap2         -------------------- -------------------- -------------------- -------------------- --------------------
D_viv_hap1         GAAATATCTTGATTCACT-A AAAAAATATTGCTGCCAATC ATACTTATCAATATATACAG ACCTAAACAGCAAAACCAAG GTTTATTATATGTTTTAACA
```

```
S_har             TAGGATCAGAAATAAAAATC TGACCCAAAGATTTCTGAAT CTATATAGTTCTAATTATAT CAGAGACATAAAGTAGATCC TGACAGAAGAAAGTAAAGAT
D_viv_hap2        -------------------- -------------------- -------------------- -------------------- --------------------
D_viv_hap1        TAGGATCAGAAATAAAAATC TGACCCAAAGATTTCTGAAT CTATATAGTTCTAATTATAT CAGAGACATAAAGTAGATCC TGACAGAAGAAAGTAAAGAT

S_har             TTACAGCACAAAGATATGCT TGTGGGACTTATATTCCAAG GATAACTAAAGATGAAAATC AGTCATATCTAGTTTACCCA AAAAAATCACTAACAAGGCT
D_viv_hap2        ------TACAAATATATATT TTACAAATATATAT------ -------------------- -------------------- --------------------
D_viv_hap1        TTACAGCACAAAGATATGCT TGTGGGACTTATATTCCAAG GATAACTAAAGATGAAAATC AGTCATATCTAGTTTACCCA AAAAAATCACTAACAAAGCT

S_har             TCTGGGATTTACAACACAGA ATGGGTATCATAAATTCTTT TATCCTCTTTGAGTTTTTTA TCATGATTGAACTAAATAGG AGTCTTAGGAAAAATTTCAG
D_viv_hap2        -------------------- -------------------- -------------------- -------------------- --------------------
D_viv_hap1        TCTGGGATTTACAACACAGA ATGGGTATCATAAATTCTTT TATCCTTTTTGAGTTTTTTA TCATGATTGAACTAAATAGG AGTCTTAGGGAAAATTTCAG

S_har             TTGGCTACTGATTCTGATTT CCCAACTATGACTTTCCTCA CAAATAAAATAGCCATGCCA ATCCCAAAATAATATTGGAG AAAAAGGAAGTTAAAAGCTC
D_viv_hap2        -------------------- -------------------- -------------------- -------------------- --------------------
D_viv_hap1        TTGGCTACTGATTCTGATTT CCCAACTATGACTTTCCTCA CAATTAAAGTAGCCATGCCA GTCCCAAAATAATATCGGAG GAAAAGAAAGTTAAAAGCTC

S_har             TTATGACAGGGGTCACCTTA TTTAAACCCTTCACTTTACA TATGGGGAAATTAAGGCCTA GAGAATTTATATAGCTTGTC CAAGGTCATACAGCTAATGA
D_viv_hap2        -------------------- -------------------- -------------------- -------------------- --------------------
D_viv_hap1        TTATGACAGAGGTCACCTTA TTTGAACCCTTCACTTTACA TATGGGAAAATTAAGGCCTA GAGAATTTATATAGCTTGTC CAAGGTCATACAGATAATGA

S_har             ATAATATAGGCAGGATGAAA ATCTTGATCCTTTTTCATCA AGTCCAGATTCCTCCTATTA TTAATGACTTCTTTAGACCA CAGGTCCTGAGGAGAACATC
D_viv_hap2        -------------------- -------------------- -------------------- -------------------- --------------------
D_viv_hap1        ATAATATAGGCAGGATGAAA ATCTTGGTCCTTTTTCATCA AGTCCAGATTCCTCCTATTA TTAATGACTTCTTTAGACCC CAGGTCCTGAGGAGAACATC

S_har             TCCAAGACCCTCAGAGATTC AGGCATACCAAAATATCTTC TTCTCCCCAAAGTACCACTT TCTCTTAATAGATCTCTTGG TCCACCATGAGTCATTCCCT
D_viv_hap2        -------------------- -------------------- -------------------- -------------------- --------------------
D_viv_hap1        TCCAAGACCCTCAGAGATCC AGGCACACCAAAATGTCTTC TTCTCCCCAAAGTACCTCTT TCTCTTAATAGATCTCTTGG TCCACCATGAGTCATTCCCT

S_har             TGGAAGAAGATTTTGAGATT TCTATTAAAACCCAAGGGTA GTGTAGCACAGTATAAATTG CCTCAGGCTGGAATCAAAAA ACCTGGGTTCTGCTCCCCTC
D_viv_hap2        -------------------- -------------------- -------------------- -------------------- --------------------
D_viv_hap1        TGGAAGAAGATTTTGAGATT TCTATTAAAACCCAAGGGTA ATTTAGCACAGTATAAATTG CCTCAGGCTGGAATCAAAAA ACCTGGGTTCTGCTCCCCTC

S_har             TTGTATGAGTTCCATTTTGG AGGCTTCAGTTTCCAAATCT ATAATACAAGAGGGTACTTT CAGAATTGATGGCCTATGAT TCTTCCCAACTCCCCCACTA
D_viv_hap2        -------------------- ---------TTTACAAATAT ATA----------------- -------------------- --------------------
D_viv_hap1        TTGTATGAGTTCCATTTTGG AGGCTTCATTTTCCAAATCT ATAATACAAGAGGGTACTTT CAGAATTGATGGCCTATGAT TCTTCCCAACTCCCCCACTA

S_har             TTTTAAAAAAAAAATTTAATT TAAAGTTTTGAGTTATAAAT TCTACATTTCCCCTCTTCCC TCCCTGAAAGAGTAAGCAAT CAGATATAGATTATACAAGT
D_viv_hap2        -------------------- -------------------- -------------------- -------------------- --------------------
D_viv_hap1        TTTTAAAAAAAATTTTAATT TAAAGTTTTGAGTTATAAAT TCTACATTTCCCCTCTTCCC TCCCTGAAAGAGTAAGCAAT CGGATATAGATTATACAAGT

S_har             GCAATTATATAAAGCTTTTA CATATTAGTTATTTTGTAAA AGAAGACCTG-AAAAAAATG AAAGAAAAATAGCATGGTTT AGTCTGGATTCAATCAAAGA
D_viv_hap2        -------------------- ---------TATTTTGTAAA AGAAGACCTGAAAAAAAATG AAAGAAAAATAGCATGATTT AGTCTGGATTCAATCAAAGA
D_viv_hap1        GCAATTATATAAAGCTTTTA CATATTAGTTATTTTGTAAA AGAAGACCTGAAAAAAAATG AAAGAAAAATAGCATGATTT AGTCTGGATTCAATCAAAGA

S_har             ACTGTATCTGTTCTTTCTCT GGAGGCAGAAAACATGCCTC ATCATTGTTTTAGGGATTGT ---CTTGGATAACTATATTT CTGAAAATAGCTAAATCATT
D_viv_hap2        ACTGTATCTGTTCTTTCTCT GGAGGCAGAAAACATGCCTC ATCATTGTTTTAGTGATTGT GGATTTGGATAACTATATTT CTGAAAATAGCTAAATCATT
D_viv_hap1        ACTGTATCTGTTCTTTCTCT GGAGGCAGAAAACATGCCTC ATCATTGTTTTAGTGATTGT GGATTTGGATAACTATATTT CTGAAAATAGCTAAATCATT

S_har             CACAATTCTTCCTTGAATGA TTTTGCTATTACTAAGTAT- AGGTTCTCCTGGTTCTGTTC TCTTCACTATGCATCAGGTT TTTCTGAAATCATCCTGCTT
D_viv_hap2        CACAATTCTTCCTTGAATGA TTTTGCTATTACTAAGTATA AGGTTCTCCTGGTTCTATTC TCTTCACTATGCATCAGGTT TTTCTGAAATCATCCTGCTT
D_viv_hap1        CACAATTCTTCCTTGAATGA TTTTGCTATTACTAAGTATA AGGTTCTCCTGGTTCTATTC TCTTCACTATGCATCAGGTT TTTCTGAAATCATCCTGCTT

S_har             GTCATTTCTTACAGCATAAT AACATTTTATTACAATCATA CAGATGTTACTGACTACTGT TTTTAGCTGTTTGACCTTGA GCAAATTACCTGACCTCTCT
D_viv_hap2        GTCATTTCTTACAGCACAAT AACATTCTATTACAATCATA CAGATATTACTGACTACTGT TTTTAGCTGTTTGACCTTGA ACAAATTACCTGACCTCTCT
D_viv_hap1        GTCATTTCTTACAGCACAAT AACATTCTATTACAATCATA CAGATATTACTGACTACTGT TTTTAGCTGTTTGACCTTGA ACAAATTACCTGACCTCTCT

S_har             GTTCAAGAATGGGTGTAATC ATTTTTCAAAATACCTCCTT CATTTGGACATTGTAAGGGA AGCACTCTAAATCTTAAAGT CCCATTATGGAAATATAATG
D_viv_hap2        GTTCAAGAATGGGTGTAATC ATTTTTCAAAATACCTCCTT CATTTGGACATTGTAAGGGA AGCACTCTAAATCTTAAAGT CCCATTATGGAAATATAATG
D_viv_hap1        GTTCAAGAATGGGTGTAATC ATTTTTCAAAATACCTCCTT CATTTGGACATTGTAAGGGA AGCACTCTAAATCTTAAAGT CCCATTATGGAAATATAATG

S_har             ATAATAATATATAATAAACA AACAAACAAATAGATAGATA GATAGATAAAAGCAAAGACC GAGGCTGCTACCTGGGAAAA ACCACCATGGGGGCCAGGCT
D_viv_hap2        ATAATAATATATAATAAACA AACAAACAA--------ATA GAGAGATAAAAGCAAAGACC GAGGCTGCTACCTGGGAAAA ACCACCATGGGGGCCAGGCT
D_viv_hap1        ATAATAATATATAATAAACA AACAAACAA--------ATA GAGAGATAAAAGCAAAGACC GAGGCTGCTACCTGGGAAAA ACCACCATGGGGGCCAGGCT

S_har             AAGTCAGAAATGTTGGGGTT CAGGGGGTGGATCGTGGGAG TAAGTGGTGGGGAGGATATC TACTGAGGGAACCAAGAGCA GGTAAGCACAAGGCTTACTG
D_viv_hap2        GAGTCAGAGATGTTGGGGTA CAGGGGGTGGATTGTGGGAG TAAGTGGTGGGGAGGATATC TACTGAGAGAACCAAGAGCA GGTAAGCCCAAGGCTTACTG
D_viv_hap1        GAGTCAGAGATGTTGGGGTA CAGGGGGTGGATTGTGGGAG TAAGTGGTGGGGAGGATATC TACTGAGAGAACCAAGAGCA GGTAAGCCCAAGGCTTACTG

S_har             TGGTAGAGAACGGGGGACTC CGAAGTCATTGGTTACCTCC TTTTTCTCAGAAAAAACCTG TGGTGAAG------TTATCT GCATCCACCTGTGCAGCCAC
D_viv_hap2        TGGTAAAGAACGGGGGACTT TGAAGTCATTGGTTACCTCC TTTTTCTCAGAAAAAAGCTG TGGTGAAGAAGTCCTCATCT GCATCCACCTGTGCAGCCAC
D_viv_hap1        TGGTAAAGAACGGGGGACTT TGAAGTCATTGGTTACCTCC TTTTTCTCAGAAAAAAGCTG TGGTGAAGAAGTCCTCATCT GCATCCACCTGTGCAGCCAC

S_har             AGGGGCCTTCTGCCAGCCTC AGACCATTTCCTGCTGCAAC AAGTGTGATACGTGCCACTG TCGCTTCTTCGGAAGTGTCT GCTTCTGCCGCCCGTTCCTG
D_viv_hap2        AGGGGCCTTCTGCCAGCCTC AGACCATTTCCTGCTGCAAC AAGTGTGATACGTGCCACTG TCGCTTCTTCGGAAGTGTCT GCTTCTGCCGCCAGTTCCTG
D_viv_hap1        AGGGGCCTTCTGCCAGCCTC AGACCATTTCCTGCTGCAAC AAGTGTGATACGTGCCACTG TCGCTTCTTCGGAAGTGTCT GCTTCTGCCGCCAGTTCCTG

S_har             CGGAAATGCTGAACGCCAGC CTCTAAGTGCTCCAAGA--G GGGGAGCCTGGGGACCCCTC CTCCCGTCACCTCCAAGGCT TCGCCCAGGGTTCCGCCCTG
D_viv_hap2        CGGAAATGCTGAACGCCAGC CTCTGAGTGCTCTAAGAGGG GGGGAGCCTGTGGACCCCTC CTCCTGTCACCTGCAAGGCT TTGCCCAGGGTTCTGCCCTG
D_viv_hap1        CGGAAATGCTGAACGCCAGC CTCTGAGTGCTCTAAGAGGG GGGGAGCCTGTGGACCCCTC CTCCTGTCACCTGCAAGGCT TTGCCCAGGGTTCTGCCCTG

S_har             ACCTGCTCAAGTAGAATTAC TTAGTGCCAGAAACCCAAAT ATGCCTCCCTCTGCTCATTT CTAAGTGACTTTCCCCGGAC ATAGCTGCTGGCTATGTCCC
D_viv_hap2        ACCTGCTCAAGTAGAATTAC TTAGTGCCAGAAACCCAAAT ATGCCTCCCTCTGCTCATTT CTAAGTGACTTTCCCTGGAC ATAGCTGCTGGCTATGTCCC
D_viv_hap1        ACCTGCTCAAGTAGAATTAC TTAGTGCCAGAAACCCAAAT ATGCCTCCCTCTGCTCATTT CTAAGTGACTTTCCCTGGAC ATAGCTGGTGGCTATGTCCC

S_har             C-TCTGGAGCCCTTGAATTC AAACTGAGAAGTCCCCAAGA TGGCAGGGAAAGTAGAGGTA ATCCTGCCCCAAATTGTGAC AGGAGAGATATCTGATTTCA
D_viv_hap2        CTTTTAGGGCCCTTGAATTC AAACTGAGAAGTCCCCAAGA TGGCAGGGAAATTAGAGGTA ATCCTGCCCCAAATTGTGAC AGGAGAAATGTCTGATTTCA
D_viv_hap1        CTTTTAGGGCCCTTGAATTC AAACTGAGAAGTCCCCAAGA TGGCAGGGAAATTAGAGGTA ATCCTGCCCCAAATTGTGAC AGGAGAAATGTCTGATTTCA

S_har             GGCCAGATGTCTGTGCCCCT CATTCCCTTCACCCCTCCTC CTCCCCCTCCCCCCAAATTC TTGTTACTTAGGATTGGGTA TTTTA
D_viv_hap2        GGCTAGATGTCTGTGCCCTT CATTCCCTTCACCCCTCCTC CTCCCCTTCCCCCCAAATTC TTGTTACTTAGGATTGGACA TTTTA
D_viv_hap1        GGCTAGATGTCTGTGCCCTT CATTCCCTTCACCCCTCCTC CTCCCCTTCCCCCCAAATTC TTGTTACTTAGGATTGGACA TTTTA

;
END;
```

### data S3: Alignment of agreodont MC1R amino acid sequences

M_lag – Greater bilby (*Macrotis lagotis*)
N_typ – Marsupial mole (*Notoryctes typhlops*)
T_cyn – Thylacine (*Thylacinus cynocephalus*)
M_fas – Numbat (*Myrmecobius fasciatus*)
S_cra – Fat-tailed dunnart (*Sminthopsis crassicaudata*)
P_tap – Brush-tailed phascogale (*Phascogale tapoatafa*)
A_fla – Yellow-footed antechinus (*Antechinus flavipes*)
A_stu – Brown antechinus (*Antechinus stuartii*)
D_viv_hap1 – Eastern quoll (*Dasyurus viverrinus*) haplotype 1
D_viv_hap2 – Eastern quoll (*Dasyurus viverrinus*) haplotype 2
S_har – Tasmanian devil (*Sarcophilus harrisii*)

Nonsense mutation
Sequence ablated by nonsense mutation

```
#NEXUS


BEGIN DATA;
 DIMENSIONS NTAX=11 NCHAR=316;
 FORMAT DATATYPE=PROTEIN INTERLEAVE MISSING=-;
[Name: M_lag           Len:   316  Check:     0]
[Name: N_typ           Len:   316  Check:     0]
[Name: M_fas           Len:   316  Check:     0]
[Name: S_cras          Len:   316  Check:     0]
[Name: P_tap           Len:   316  Check:     0]
[Name: A_fla           Len:   316  Check:     0]
[Name: A_stu           Len:   316  Check:     0]
[Name: D_viv_hap1      Len:   316  Check:     0]
[Name: D_viv_hap2      Len:   316  Check:     0]
[Name: S_har           Len:   316  Check:     0]
[Name: T_cyn           Len:   316  Check:     0]

MATRIX
M_lag                 MPMPGQQKRLFNSLNATPPD TLHQVVPTNQTDIPCQGLFI PDELFLTLGLLSLVENVMVM VAIIKNRNLHSPMYYFVCCL ALSDLLVSVSNLLETLVMLL
N_typ                 MPMPGQQKRLFNSLNFTSPD TRHQVVPTNQTDIPCQGLFI PDELFLSLGLLSLVENVIVV AAIIKNRNLHSPMYYFVCCL ALSDLLVSVSNLLETLVMLL
M_fas                 MPMPGQQKRLFNSLNATSPA TLHQVVPTNQTAIPCQVLLI PEELFLTLGLLSLVENVMVV VAIVKNRNLHSPMYYFVCCL ALSDLLVSVSNLLETLVVLL
S_cras                MPMPGQQKRLFNTLNATSPG PLHRVVPANRTEFPCQVLFV PDELFLTLGLLSLLENVMVV VAIVKNRNLHAPMYYFVCCL ALSDLLVSVSNLLETVVGLL
P_tap                 MPMPGQQKRLFNSLNATAPD TLHQVVPANRTAFPCQVLFV PEELFLTLGLLSLLENIMVV VAIAKNRNLHAPMYYFVCCL ALSDLLVSVSSLLETVVVLL
A_fla                 MPMPGQQKRLFNSLNGTAPD TLHQVVPANRTDFPCQVLFV PDELFLTLGLLSLLENVMVV VAIAKNRNLHAPMYYFVCCL ALSDLLVSVSSLLETVVVLL
A_stu                 MPMPGQQKRLFNSLNGTAPD TLHQVVPANRTDFPCQVLFV PDELFLTLGLLSLLENVMVV VAIAKNRNLHAPMYYFVCCL ALSDLLVSVSSLLETVVVLL
D_viv_hap1            MPMPGQQKRLFNSLNATSPD ALHQVVPANRTDFPCQVLFV PNALFLTLGLLSLLENVMVV VAIAKNRNLHAPMYYFVCCL ALSDLLVSVSNLLETVVVLL
D_viv_hap2            MPMPGQQKRLFNSLNATSPD ALHQVVPANRTDFPCQVLFV PNALFLTLGLLSLLENVMVV VAIAKNRNLHAPMYYFVCCL ALSDLLVSVSNLLETVVVLL
S_har                 MPMPGQQKRLFNSLNATSPD ALHQVVPANRTDFPCQVLFV PNALFLTLGLLSLLENIMVV VAITKNRNLHAPMYYFVCCL ALSDLLVSVSNLLETVVVLL
T_cyn                 MPMPGQQKRLFNSLNATAPD AFHQVAPANRTEILCQVLFI PDELFLTLGLLSLVENVMVV VAILKNHNLHSPMYYFVCCL ALSDLLVSVSNLLETLVVLP

M_lag                 LEKGVLVIQASTVQQLDNVI DVLICGSMMSSISFLGAIAV DRYISIFYALRYHSIVTPCR AQGVLAGIWVSSALLGTLFI SYYNHVAVLLCLIGFFLSML
N_typ                 LEKGVLVIQAPMMQQLDNVI DVLICGSMMSSISFLGAIAV DRYISIFYALRYHSIVTPC- AQAALAGIWVSSALFGTLFI SYYNHVVVLLCLIGFFLSML
M_fas                 LERRLLAIQAATVQQLNNVI DVLICGSMMSSISFLGAIAV DRYISIFYALRYHSIVTPCR AQGVLAGIWVTSALSGALSI SYYNHVAVLLCLIGFFLSML
S_cras                LERRVLAIRAPAVQQLNNVI DVLICGSMMSSISFLGAIAV DRYISIFYALRYRSIVTSCR AQGVLAGIWVSSALSGALFI SYYNHVAVLLCLIGFFLSTL
P_tap                 LERGLLAMRAPAAQQLNNVI DVLICGSMMSSISFLGAIAV DRYISIFYALRYRSIVTPCR AQGVLAGIWVSSALSGALFI AYYNHVAVLLCLISFFLSML
A_fla                 LERGLLAMRAPAAQQLNNVI DVLICGSMMSSISFLGAIAV DRYISIFYALRYRSIVTPGR AQGVLAGIWVASALSGALFI AYYNHVAVLLCLIGFFLSML
A_stu                 LERGLLAMRAPAAQQLNNVI DVLICGSMMSSISFLGAIAV DRYISIFYALRYRSIVTPGR AQGVLAGIWVASALSGALFI AYYNHVAVLLCLIGFFLSML
D_viv_hap1            LERGLLAIQAPAVQRLNNVI DVLICGSMMSSISFLGAIAV DRYISIFYALRYRSIVTPCR AQGVLAGIWVSSALSGILFI SYYNHVVVLLCLIGFFLSML
D_viv_hap2            LERGLLAIQAPAVQRLNNVI DVLICGSMMSSISFLGAIAV DRYISIFYALRYRSIVTPCR AQGVLAGIWVSSALSGTLFI SYYNHVVVLLCLIGFFLSML
S_har                 LERGLLAIQAPVVQRLNNVI DVLICGSMMSSISFLGAIAV DRYISIFYALRYRSIVTPCR AQGVLAGIWVSSALSGTLFI SYYNHVAVLLCLIGFFLSML
T_cyn                 LERGVLAIQAPTMQQLDNVI DVLICGSMMSSISFLGAIAV DRYISIFYALRYHSIVTPCR AQGVLAGIWVSSALSGALFI SYYNHIAVLLCLIGFFLSML

M_lag                 GLMVVLYIHMFIQACQHARR IARLHKRYTIHQLSTLKGAV TLTILLGIFFLCWAPFFLHL TLIVLCPKHPTCTCYFQNFN LFLILILCNSVIDPLIYAFR
N_typ                 GLMVVLYIHMFIQACQHARR IARLHKRYTIHQMSTLKGAV TLTILLGIFFLCWTPFFLHL TLIVLCPKHPTCSCYFQNFN LFLILIICNSVIDPLIYAFR
M_fas                 GLMVVLYIHMFVRACQHARR IAQLHKRYPIHQLSTLKGAV TLTILLGIFFLCWAPFFLHL TLIVLCPKHPTCVCYFQNFN LFLILIICNSIIDPLIYAFR
S_cras                GLMVVLYIHMFIRACQHARR IARLHKRYPVHQLSPLRGAV TLTILLGIFFLCWAPFFLHL TLIVLCPKHPMCSCYFQNFN LFLILIICNSVIDPLIYAFR
P_tap                 GLMVVLYIHMFIRACQHARR IAQLHKRYPVHQLSTLRGAV TLTILLGIFFLCWAPFFLHL TLIVLCPKHPTCSCYFQNFN LFLILIICNSVIDPLIYAFR
A_fla                 GLMVVLYIHMFVRACQHARR ISRLHKRHPVHQLSTLRGAV TLTILLGIFFLCWAPFFLHL TLIVLCPEHPTCSCYFQNFN LFLILIICNSVIDPLIYAFR
A_stu                 GLMVVLYIHMFIRACQHARR ISRLHKRYPVHQLSTLRGAV TLTILLGIFFLCWAPFFLHL TLIVLCPKHPTCSCYFQNFN LFLILIICNSVIDPLIYAFR
D_viv_hap1            GLMVVLYIHMFIRACQHARR IARLHKRYPVHQLSTLRGAV TLTILLGIFFLCWAPFFLHL TLIVLCPKHPTCSCYFQNFN LFLILIICNSVIDPLIYAFR
D_viv_hap2            GLMVVLYIHMFIRACQHARR IARLHKRYPVHQLSTLRGAV TLTILLGIFFLCWAPFFLHL TLIVLCPKHPTCSCYFQNFN LFLILIICNSVIDPLIYAFR
S_har                 GLMVVLYIHMFIRACQHARR ITRLHKRYPVHQLSILRGAV TLTILLGIFFLCWAPFFLHL TLIVLCPKHPTCSCYFQNFN LFLILIICNSVIDPLIYAFR
T_cyn                 GLMVVLYIHMFIQACQHARR IAQLHKGYPIHHLSTLKGAV TLTILLGIFFLCWAPFFLHL TLIVLCPKHPTCSCYFQNFN LFLILIICNSVIDPLIYAFR

M_lag                 SQELCKTLKEVILCSW
N_typ                 SQELRKTLKEVILCSW
M_fas                 SQELRKTLKEVVLCSW
S_cras                SQELRKTLKEVILCSW
P_tap                 SQELRKTLKEVILCSW
A_fla                 SQELRKTLKEVILCSW
A_stu                 SQELRKTLKEVILCSW
D_viv_hap1            SQELRKTLKEVILCSW
D_viv_hap2            SQELRKTLKEVILCSW
S_har                 SQELRKTLKEVILCSW
T_cyn                 SQELRKTLKEMILCSW

;
END;
```

```
#NEXUS


BEGIN DATA;
 DIMENSIONS NTAX=11 NCHAR=951;
 FORMAT DATATYPE=DNA INTERLEAVE MISSING=-;
[Name: M_lag            Len:   951  Check:     0]
[Name: N_typ            Len:   951  Check:     0]
[Name: T_cyn            Len:   951  Check:     0]
[Name: M_fas            Len:   951  Check:     0]
[Name: S_cras           Len:   951  Check:     0]
[Name: P_tap            Len:   951  Check:     0]
[Name: A_fla            Len:   951  Check:     0]
[Name: A_stu            Len:   951  Check:     0]
[Name: D_viv_hap1       Len:   951  Check:     0]
[Name: D_viv_hap2       Len:   951  Check:     0]
[Name: S_har            Len:   951  Check:     0]

MATRIX
M_lag                ATGCCGATGCCAGGTCAACA GAAGAGATTATTTAACTCTC TGAATGCCACTCCCCCAGAT ACCCTCCATCAAGTTGTCCC CACCAACCAGACAGATATTC
N_typ                ATGCCAATGCCAGGTCAACA GAAGAGATTGTTTAACTCTC TAAACTTCACTTCCCCAGAT ACGCGCCATCAAGTTGTCCC CACCAACCAGACAGATATTC
T_cyn                ATGCCGATGCCGGGTCAACA GAAGAGATTGTTTAACTCTC TGAACGCCACTGCCCCGGAC GCCTTCCATCAAGTTGCCCC CGCCAACCGGACGGAGATCC
M_fas                ATGCCGATGCCGGGTCAACA GAAGAGATTGTTTAACTCCC TGAACGCCACTTCCCCGGCC ACCCTCCATCAAGTTGTCCC CACCAACCAGACGGCCATCC
S_cras               ATGCCGATGCCGGGTCAACA GAAGCGGTTGTTTAACACTC TGAACGCCACTTCCCCGGGC CCCCTCCATCGGGTCGTCCC CGCCAACCGGACGGAGTTCC
P_tap                ATGCCGATGCCGGGTCAACA GAAGCGGCTGTTTAACTCTC TGAACGCCACTGCCCCGGAC ACCCTCCACCAGGTCGTCCC CGCCAACCGGACGGCCTTCC
A_fla                ATGCCGATGCCGGGTCAACA GAAGCGGCTGTTTAACTCTC TGAACGGCACTGCCCCGGAC ACCCTCCACCAGGTCGTCCC CGCCAACCGGACGGACTTCC
A_stu                ATGCCGATGCCGGGTCAACA GAAGCGGCTGTTTAACTCTC TGAACGGCACTGCCCCGGAC ACCCTCCACCAGGTCGTCCC CGCCAACCGGACGGACTTCC
D_viv_hap1           ATGCCGATGCCGGGTCAACA GAAGCGGCTGTTTAACTCTC TGAACGCCACTTCCCCAGAC GCCCTTCACCAGGTGGTCCC CGCCAACCGGACGGACTTCC
D_viv_hap2           ATGCCGATGCCGGGTCAACA GAAGCGGCTGTTTAACTCTC TGAACGCCACTTCCCCAGAC GCCCTTCACCAGGTGGTCCC CGCCAACCGGACGGACTTCC
S_har                ATGCCGATGCCGGGTCAACA GAAGCGGCTGTTTAACTCTC TGAACGCCACTTCCCCGGAC GCCCTCCACCAGGTGGTCCC CGCCAACCGGACGGACTTCC

M_lag                CATGCCAGGGGCTCTTCATT CCAGATGAGTTGTTCTTGAC CCTGGGGCTTCTAAGCCTGG TGGAGAATGTCATGGTGATG GTGGCCATCATCAAGAACCG
N_typ                CTTGCCAGGGGCTCTTCATT CCAGATGAGTTGTTCTTGAG CTTGGGGCTCCTGAGCCTGG TGGAGAATGTCATAGTAGTG GCGGCCATCATCAAGAACCG
T_cyn                TGTGCCAGGTGCTGTTCATT CCGGATGAGTTATTCTTGAC CCTGGGGCTGCTGAGCCTGG TGGAGAATGTCATGGTGGTG GTGGCCATCCTCAAGAACCA
M_fas                CGTGCCAGGTGCTGCTCATT CCCGAGGAGCTCTTCTTGAC CCTGGGGCTGCTGAGCCTGG TGGAGAACGTCATGGTGGTG GTGGCCATCGTCAAGAACCG
S_cras               CGTGCCAGGTGCTCTTCGTG CCCGACGAGCTCTTCTTGAC CCTAGGGCTGCTGAGCCTGC TGGAGAACGTCATGGTGGTG GTGGCCATCGTCAAGAACCG
P_tap                CGTGCCAGGTGCTCTTCGTG CCCGAGGAGCTCTTCCTGAC ACTGGGGCTGCTGAGCCTGC TGGAGAACATCATGGTGGTG GTGGCCATCGCCAAGAACCG
A_fla                CGTGCCAGGTGCTCTTCGTG CCCGACGAGCTCTTCCTGAC GCTGGGGCTGCTGAGCCTGC TGGAGAACGTCATGGTGGTG GTGGCCATCGCCAAGAACCG
A_stu                CGTGCCAGGTGCTCTTCGTG CCCGACGAGCTCTTCCTGAC GCTGGGGCTGCTGAGCCTGC TGGAGAACGTCATGGTGGTG GTGGCCATCGCCAAGAACCG
D_viv_hap1           CGTGCCAGGTGCTCTTTGTG CCCAATGCGCTCTTCCTGAC GCTGGGGCTGCTGAGCCTGC TGGAGAACGTCATGGTGGTG GTGGCCATCGCCAAGAACCG
D_viv_hap2           CGTGCCAGGTGCTCTTTGTG CCCAATGCGCTCTTCCTGAC GCTGGGGCTGCTGAGCCTGC TGGAGAACGTCATGGTGGTG GTGGCCATCGCCAAGAACCG
S_har                CGTGCCAGGTGCTCTTTGTG CCCAATGCGCTTTTCCTGAC ACTGGGGCTGCTGAGCCTGC TGGAGAACATCATGGTGGTG GTGGCCATCACCAAGAACCG

M_lag                CAACCTGCATTCACCCATGT ATTATTTTGTCTGCTGCCTG GCTCTATCAGACCTTCTGGT GAGTGTCAGCAACCTACTGG AGACCTTGGTGATGCTACTG
N_typ                CAACCTGCATTCACCCATGT ACTATTTTGTCTGCTGCCTG GCTCTGTCAGACCTTCTGGT GAGCGTCAGCAATCTGCTGG AGACCTTGGTCATGCTACTC
T_cyn                CAACCTGCATTCGCCCATGT ACTATTTTGTCTGCTGCCTG GCTCTGTCAGACCTTCTGGT GAGCGTCAGCAACCTGCTGG AGACCTTGGTGGTGCTGCCA
M_fas                CAACCTGCATTCGCCCATGT ACTACTTTGTCTGCTGCCTG GCGCTCTCAGACCTGCTGGT GAGCGTCAGCAACCTGCTGG AGACCTTGGTGGTGCTGCTG
S_cras               CAACCTGCACGCCCCCATGT ACTATTTTGTCTGCTGCCTG GCTCTGTCGGACCTGCTGGT GAGTGTCAGCAACCTGCTGG AGACCGTGGTGGGGCTGCTG
P_tap                CAACCTGCACGCCCCCATGT ACTATTTTGTCTGCTGCCTG GCTCTGTCGGACCTTCTGGT GAGTGTCAGCAGCCTGCTGG AGACCGTGGTGGTGCTGCTG
A_fla                CAACCTGCACGCCCCCATGT ACTATTTTGTCTGCTGCCTG GCTCTGTCGGACCTTCTGGT GAGTGTCAGCAGCCTGCTGG AGACCGTGGTGGTGCTGCTG
A_stu                CAACCTGCACGCCCCCATGT ACTATTTTGTCTGCTGCCTG GCTCTGTCGGACCTTCTGGT GAGTGTCAGCAGCCTGCTGG AGACCGTGGTGGTGCTGCTG
D_viv_hap1           CAACCTGCACGCCCCCATGT ACTATTTTGTCTGCTGCCTG GCTCTGTCGGACCTTCTGGT GAGTGTCAGCAACCTGCTGG AGACCGTGGTGGTGCTGCTG
D_viv_hap2           CAACCTGCACGCCCCCATGT ACTATTTTGTCTGCTGCCTG GCTCTGTCGGACCTTCTGGT GAGTGTCAGCAACCTGCTGG AGACCGTGGTGGTGCTGCTG
S_har                CAACCTGCACGCCCCCATGT ACTATTTTGTCTGCTGCCTG GCTCTGTCGGACCTTCTGGT GAGTGTCAGCAACCTGCTGG AGACCGTGGTGGTGCTGCTG

M_lag                CTGGAGAAAGGGGTTCTGGT GATCCAGGCATCAACGGTAC AACAGCTTGACAATGTCATT GATGTGTTGATCTGTGGGTC CATGATGTCTTCAATTTCCT
N_typ                CTGGAGAAAGGGGTGCTGGT AATCCAGGCACCTATGATGC AACAACTTGACAATGTCATT GATGTGTTGATCTGTGGTTC CATGATGTCCTCCATTTCCT
T_cyn                CTGGAGAGAGGGGTGCTGGC CATCCAGGCGCCCACCATGC AACAGCTCGACAACGTCATC GACGTCTTGATCTGCGGCTC CATGATGTCCTCCATTTCCT
M_fas                CTGGAGAGGAGGCTGCTGGC CATCCAGGCGGCCACGGTGC AACAGCTAAACAACGTCATT GACGTGCTGATCTGTGGCTC CATGATGTCCTCCATTTCCT
S_cras               CTGGAGAGGAGGGTGCTGGC CATCCGGGCGCCCGCAGTGC AGCAGCTAAACAATGTCATT GATGTGTTGATCTGTGGTTC CATGATGTCCTCCATTTCCT
P_tap                CTGGAGAGGGGGCTGCTGGC CATGCGGGCGCCCGCGGCGC AGCAGCTAAACAATGTCATT GATGTGTTGATCTGTGGTTC CATGATGTCCTCCATTTCCT
A_fla                CTGGAGAGGGGGCTGCTGGC CATGCGGGCGCCCGCGGCGC AGCAGCTAAACAATGTCATT GATGTGTTGATCTGTGGTTC CATGATGTCCTCCATTTCCT
A_stu                CTGGAGAGGGGGCTGCTGGC TATGCGGGCGCCCGCGGCGC AGCAGCTAAACAATGTCATT GATGTGTTGATCTGTGGTTC CATGATGTCCTCCATTTCCT
D_viv_hap1           CTGGAGAGGGGGCTGCTGGC CATCCAGGCGCCCGCGGTGC AGCGGCTAAACAATGTCATT GATGTGTTGATCTGTGGTTC CATGATGTCCTCCATTTCCT
D_viv_hap2           CTGGAGAGGGGGCTGCTGGC CATCCAGGCGCCCGCGGTGC AGCGGCTAAACAATGTCATT GATGTGTTGATCTGTGGTTC CATGATGTCCTCCATTTCCT
S_har                CTGGAGAGGGGGCTGCTGGC CATCCAGGCGCCCGTGGTGC AGCGGCTAAACAATGTCATT GATGTGTTGATCTGTGGTTC CATGATGTCCTCCATTTCCT

M_lag                TCCTAGGAGCCATTGCTGTT GACCGCTACATTAGTATCTT CTATGCCCTTCGCTACCACA GCATAGTCACTCCTTGCCGT GCTCAGGGAGTCCTTGCTGG
N_typ                TCCTAGGAGCTATTGCTGTT GACCGCTATATCAGTATCTT CTATGCCCTGCGCTACCACA GCATTGTCACCCCTTGTTGA GCTCAGGCAGCCCTTGCTGG
T_cyn                TCCTGGGAGCCATCGCCGTC GATCGCTACATCAGTATCTT CTATGCCCTGCGCTACCACA GCATCGTCACTCCCTGTCGC GCTCAGGGAGTCCTTGCTGG
M_fas                TCCTAGGAGCCATCGCCGTG GACCGCTACATCAGCATCTT CTACGCCCTGCGCTACCACA GCATTGTCACCCCCTGCCGA GCTCAGGGCGTCCTGGCCGG
S_cras               TCCTAGGAGCCATTGCTGTG GACCGCTACATCAGCATCTT CTATGCCCTGCGCTACCGCA GCATCGTCACGTCCTGCAGG GCCCAGGGGGTCCTCGCCGG
P_tap                TCCTAGGAGCCATCGCCGTG GACCGCTACATCAGCATCTT CTACGCCCTGCGCTACCGCA GCATCGTCACGCCCTGCCGG GCCCAGGGCGTCCTCGCCGG
A_fla                TCCTAGGAGCCATCGCCGTG GACCGCTACATCAGCATCTT CTACGCCCTGCGCTACCGCA GCATCGTCACGCCCGGCCGG GCCCAGGGCGTCCTCGCCGG
A_stu                TCCTAGGAGCCATCGCCGTG GACCGCTACATCAGCATCTT CTACGCCCTGCGCTACCGCA GCATCGTCACGCCCGGCCGG GCCCAGGGCGTCCTCGCCGG
D_viv_hap1           TCCTAGGAGCCATCGCCGTG GACCGCTACATCAGCATCTT CTATGCCCTGCGCTACCGCA GCATCGTCACGCCCTGCCGG GCCCAGGGGGTGCTCGCCGG
D_viv_hap2           TCCTAGGAGCCATCGCCGTG GACCGCTACATCAGCATCTT CTATGCCCTGCGCTACCGCA GCATCGTCACGCCCTGCCGG GCCCAGGGGGTGCTCGCCGG
S_har                TCCTAGGAGCCATCGCCGTG GACCGCTACATCAGTATCTT CTATGCCCTGCGCTACCGCA GCATCGTCACGCCCTGCCGG GCCCAGGGGGTGCTCGCCGG

M_lag                CATCTGGGTGTCCAGTGCTC TCTTGGGTACCCTCTTCATC TCCTACTACAACCATGTTGC AGTCCTACTCTGTCTCATTG GCTTCTTCTTGTCCATGTTG
N_typ                CATCTGGGTGTCCAGTGCCC TCTTTGGTACCCTCTTCATC TCCTACTATAACCATGTTGT AGTCCTGCTCTGTCTCATTG GCTTTTTCTTGTCCATGTTG
T_cyn                CATCTGGGTGTCCAGCGCCC TGTCCGGCGCCCTCTTCATC TCCTATTACAACCACATCGC GGTTCTGCTCTGCCTCATTG GCTTCTTCTTGTCCATGTTG
M_fas                CATCTGGGTGACCAGTGCCC TGTCCGGTGCCCTCTCCATC TCCTATTACAACCACGTGGC GGTGCTGCTCTGCCTCATCG GCTTCTTCTTGTCCATGTTG
S_cras               CATCTGGGTGTCTAGCGCCC TATCGGGCGCCCTCTTCATC TCCTACTACAACCACGTGGC GGTCCTGCTCTGCCTCATCG GCTTCTTCTTGTCCACGCTA
P_tap                CATCTGGGTGTCCAGCGCGC TGTCGGGCGCCCTCTTCATC GCCTACTATAACCACGTGGC GGTCCTGCTCTGTCTCATCA GCTTCTTCTTGTCCATGCTG
A_fla                CATCTGGGTGGCCAGCGCGC TGTCGGGCGCCCTCTTCATC GCCTACTACAACCACGTGGC GGTCCTGCTCTGTCTCATCG GCTTCTTCTTGTCCATGCTG
A_stu                CATCTGGGTGGCCAGCGCGC TGTCGGGCGCCCTCTTCATC GCCTACTACAACCACGTGGC GGTCCTGCTCTGTCTCATCG GCTTCTTCTTGTCCATGCTG
D_viv_hap1           CATCTGGGTGTCCAGCGCCC TGTCGGGCATCCTCTTCATC TCCTACTACAACCATGTGGT GGTCCTGCTCTGCCTTATCG GCTTCTTCTTGTCCATGCTG
D_viv_hap2           CATCTGGGTGTCCAGCGCCC TGTCGGGCACCCTCTTCATC TCCTACTACAACCATGTGGT GGTCCTGCTCTGCCTTATCG GCTTCTTCTTGTCCATGCTG
S_har                CATCTGGGTGTCCAGTGCCC TGTCGGGCACCCTCTTCATC TCCTACTACAACCACGTGGC GGTCCTGCTCTGCCTTATCG GCTTCTTCTTGTCCATGCTG

M_lag                GGGCTCATGGTGGTCCTCTA CATTCACATGTTCATCCAGG CATGCCAGCATGCCAGGAGG ATTGCTCGGCTACACAAGAG ATACACCATTCACCAGCTGT
N_typ                GGGCTTATGGTGGTCCTCTA CATTCACATGTTCATCCAGG CATGCCAGCATGCCAGGAGG ATTGCTCGACTACACAAGAG ATACACCATTCACCAGATGT
T_cyn                GGGCTCATGGTGGTCCTCTA CATTCACATGTTCATCCAGG CGTGCCAGCACGCCAGGAGG ATCGCTCAGCTCCACAAGGG ATACCCCATTCACCATCTGT
M_fas                GGGCTCATGGTGGTCCTCTA CATCCACATGTTCGTCCGGG CGTGCCAGCATGCCCGGAGG ATCGCTCAGCTCCACAAGAG GTACCCCATTCACCAGCTGT
S_cras               GGGCTCATGGTGGTCCTCTA CATCCACATGTTCATCCGAG CGTGCCAGCATGCCAGAAGG ATCGCGCGGCTGCACAAGAG ATACCCCGTCCACCAGCTGT
P_tap                GGGCTCATGGTGGTCCTCTA CATCCACATGTTCATCCGGG CGTGCCAGCACGCCAGGAGG ATCGCGCAGCTGCACAAGAG ATACCCCGTCCACCAGCTGT
A_fla                GGGCTCATGGTGGTCCTCTA CATCCACATGTTCGTCCGGG CGTGCCAGCACGCCAGGAGG ATCTCGCGGCTGCACAAGAG ACACCCCGTCCACCAGCTGT
A_stu                GGGCTCATGGTGGTCCTCTA CATCCACATGTTCATCCGGG CGTGCCAGCACGCCAGGAGG ATCTCGCGGCTGCACAAGAG ATACCCCGTCCACCAGCTGT
D_viv_hap1           GGGCTCATGGTGGTCCTCTA CATCCACATGTTCATCCGGG CGTGCCAACACGCCAGGAGG ATCGCGCGGCTGCACAAGAG ATACCCTGTGCACCAGCTGT
D_viv_hap2           GGGCTCATGGTGGTCCTCTA CATCCACATGTTCATCCGGG CGTGCCAACACGCCAGGAGG ATCGCGCGGCTGCACAAGAG ATACCCTGTGCACCAGCTGT
S_har                GGGCTCATGGTGGTCCTCTA CATCCACATGTTCATCCGGG CATGCCAGCACGCCAGGAGG ATCACGCGGCTGCACAAGAG ATACCCCGTGCACCAGCTGT
```

```
M_lag           CAACCCTCAAGGGGGCTGTC ACCCTCACAATTCTGTTGGG CATCTTCTTCCTCTGTTGGG CCCCCTTTTTCCTGCATCTC ACACTAATTGTCCTCTGTCC
N_typ           CAACCCTCAAGGGGGCTGTC ACCCTCACAATCCTGCTAGG CATCTTCTTCCTCTGCTGGA CCCCTTTTTTCTTGCACCTC ACACTTATTGTCCTCTGTCC
T_cyn           CAACCCTCAAAGGGGCCGTC ACCCTCACAATCCTATTGGG CATCTTCTTCCTCTGCTGGG CCCCCTTTTTCCTGCACCTC ACGCTTATTGTCCTCTGTCC
M_fas           CGACGCTCAAAGGGGCTGTC ACCCTCACGATCCTCTTGGG CATCTTCTTTCTCTGCTGGG CCCCCTTCTTCCTGCACCTC ACGCTCATCGTCCTCTGTCC
S_cras          CCCCCCTCAGGGGGGCCGTC ACCCTCACCATCCTGCTGGG CATCTTCTTCCTCTGCTGGG CCCCCTTTTTCCTGCACCTC ACGCTTATCGTCCTTTGCCC
P_tap           CGACCCTCAGGGGGGCCGTC ACCCTCACCATCCTGCTGGG CATCTTCTTCCTCTGCTGGG CCCCCTTTTTCCTGCACCTC ACGCTCATCGTCCTCTGCCC
A_fla           CGACGCTCAGGGGGGCCGTC ACCCTCACCATCCTGCTGGG CATCTTCTTCCTCTGCTGGG CCCCCTTTTTCCTGCACCTC ACGCTCATCGTCCTCTGCCC
A_stu           CGACGCTCAGGGGGGCCGTC ACCCTCACCATCCTGCTGGG CATCTTCTTCCTCTGCTGGG CCCCCTTTTTCCTGCACCTC ACGCTCATCGTCCTCTGCCC
D_viv_hap1      CGACCCTCAGAGGGGCCGTC ACCCTCACCATCTTGCTGGG CATCTTCTTCCTCTGCTGGG CCCCCTTTTTCCTGCACCTC ACGCTCATCGTCCTCTGCCC
D_viv_hap2      CGACCCTCAGAGGGGCCGTC ACCCTCACCATCTTGCTGGG CATCTTCTTCCTCTGCTGGG CCCCCTTTTTCCTGCACCTC ACGCTCATCGTCCTCTGCCC
S_har           CGATCCTCAGAGGGGCTGTC ACCCTCACCATCCTGCTGGG CATCTTCTTCCTCTGCTGGG CCCCCTTTTTCCTGCACCTC ACGCTCATCGTCCTCTGCCC

M_lag           CAAGCATCCCACATGCACCT GCTACTTCCAGAACTTCAAC CTCTTTCTCATCCTCATCCT CTGCAACTCGGTCATTGATC CTCTCATCTATGCCTTCCGC
N_typ           CAAGCATCCCACATGCAGCT GCTACTTCCAGAACTTCAAC CTCTTTCTCATCCTCATCAT CTGCAACTCGGTCATTGACC CTCTCATCTATGCCTTCCGT
T_cyn           CAAGCATCCCACATGCAGCT GCTACTTTCAGAACTTCAAC CTCTTTCTCATCCTCATCAT CTGCAACTCAGTCATTGACC CTCTCATCTATGCCTTCCGC
M_fas           CAAGCACCCCACGTGCGTCT GCTACTTTCAGAACTTCAAC CTCTTTCTCATCCTCATCAT CTGCAACTCGATCATCGACC CTCTCATCTATGCCTTCCGC
S_cras          CAAGCATCCCATGTGCAGCT GCTACTTTCAGAACTTCAAC CTTTTCCTCATCCTCATCAT CTGCAACTCAGTCATTGACC CTCTCATCTACGCCTTCCGC
P_tap           CAAGCATCCCACGTGCAGCT GCTACTTTCAGAACTTCAAC CTCTTTCTCATCCTCATCAT CTGCAACTCGGTCATCGACC CGCTCATCTACGCCTTCCGC
A_fla           CGAGCATCCCACGTGCAGCT GCTACTTTCAGAACTTCAAC CTCTTTCTCATCCTCATCAT CTGCAACTCAGTCATCGACC CGCTCATCTACGCCTTCCGC
A_stu           CAAGCATCCCACGTGCAGCT GCTACTTTCAGAACTTCAAC CTCTTTCTCATCCTCATCAT CTGCAACTCAGTCATCGACC CGCTCATCTACGCCTTCCGC
D_viv_hap1      CAAGCATCCCACGTGCAGCT GCTACTTTCAGAACTTCAAC CTCTTTCTCATCCTCATCAT CTGCAACTCAGTCATTGACC CGCTCATCTACGCCTTCCGC
D_viv_hap2      CAAGCATCCCACGTGCAGCT GCTACTTTCAGAACTTCAAC CTCTTTCTCATCCTCATCAT CTGCAACTCAGTCATTGACC CGCTCATCTACGCCTTCCGC
S_har           CAAGCATCCCACGTGCAGCT GCTACTTTCAGAACTTCAAC CTCTTTCTCATCCTCATCAT CTGCAACTCAGTCATTGACC CGCTCATCTACGCCTTCCGC

M_lag           AGCCAAGAACTCTGCAAGAC CCTCAAGGAGGTGATTCTAT GTTCCTGGTAA
N_typ           AGCCAAGAACTCCGCAAGAC ACTCAAGGAGGTGATTCTGT GTTCCTGGTAA
T_cyn           AGCCAAGAGCTCCGCAAGAC CCTCAAGGAGATGATTCTGT GTTCCTGGTAA
M_fas           AGCCAAGAGCTCCGCAAGAC CCTCAAGGAGGTGGTTCTGT GTTCCTGGTAA
S_cras          AGCCAGGAGCTCCGCAAGAC GCTCAAGGAGGTGATTCTGT GTTCCTGGTGA
P_tap           AGCCAGGAGCTCCGCAAGAC CCTCAAGGAGGTGATCCTGT GTTCCTGGTGA
A_fla           AGCCAGGAGCTCCGCAAGAC CCTCAAGGAGGTGATCCTGT GTTCCTGGTGA
A_stu           AGCCAGGAGCTCCGCAAGAC CCTCAAGGAGGTGATCCTGT GTTCCTGGTGA
D_viv_hap1      AGCCAGGAGCTCCGCAAGAC CCTCAAGGAGGTGATTCTGT GTTCCTGGTGA
D_viv_hap2      AGCCAGGAGCTCCGCAAGAC CCTCAAGGAGGTGATTCTGT GTTCCTGGTGA
S_har           AGCCAGGAGCTCCGCAAGAC CCTCAAGGAGGTGATTCTGT GTTCCTGGTGA

;
END;
```

figure S1

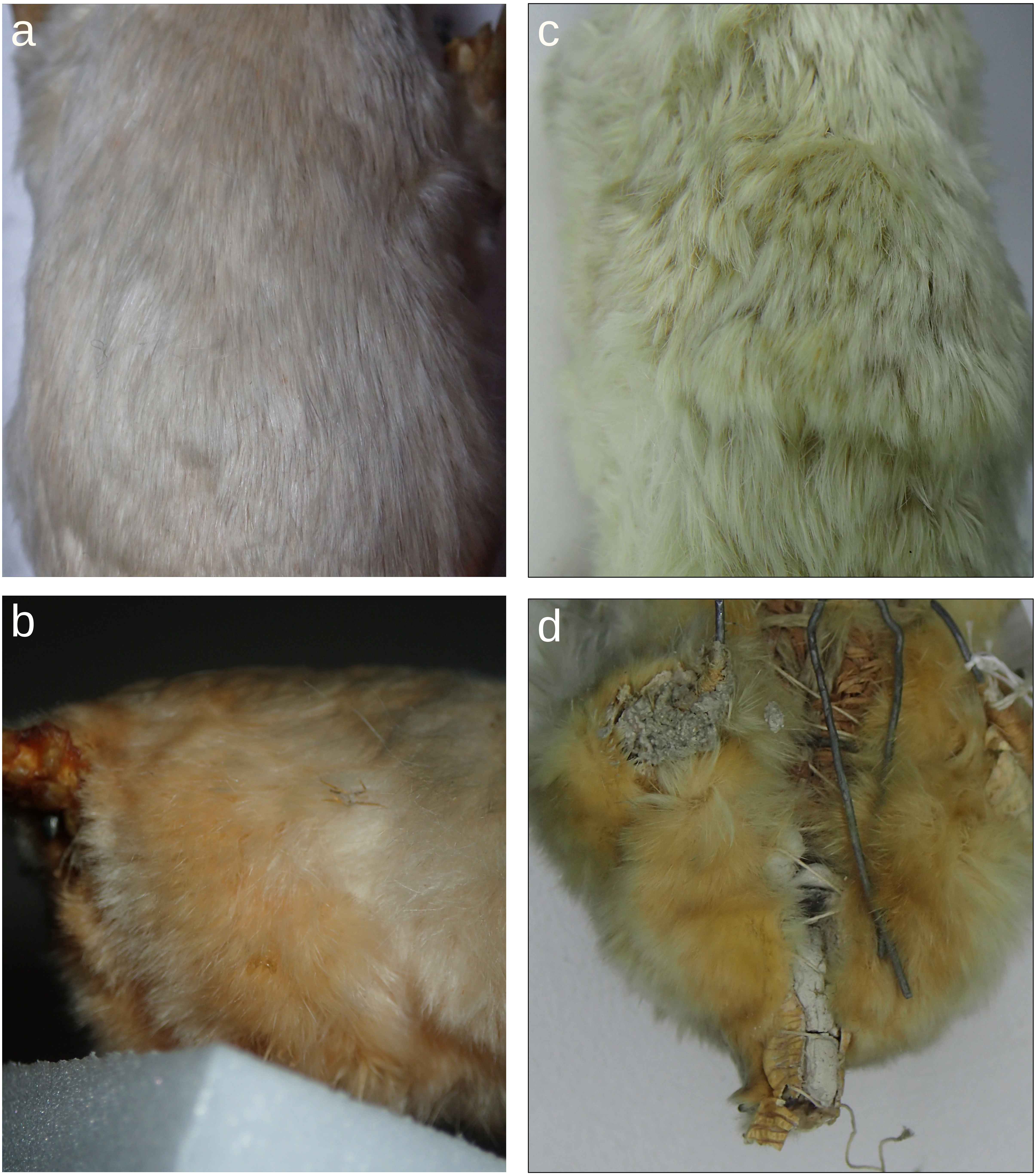


Figure S1: Photos of coat colour in marsupial moles. Dorsal fur (top row) tends to exhibit a much paler yellow colour compared to fur on the ventral posterior aspect of the limbs (bottom row), as illustrated by two mounted specimens at Melbourne Museum, C25067 (a & b) and C2901 (c & d).

**code S1: Script to plot region alignment between eastern quoll haplotypes at the ASIP locus**

```
# Load all packages
library(SVbyEye)
library(rtracklayer)
library(GenomicRanges)
library(ggplot2)
library(svglite)



# Load paf alignment, read it to a table
paf.file <- "haplotype_minimap2_aln.paf"
paf.table <- readPaf(paf.file = paf.file, include.paf.tags = TRUE, restrict.paf.tags
= "cg")

# Subset paf alignment to asip locus

paf.subset <- subsetPafAlignments(paf.table = paf.table, target.region =
"target:422951000-422960000")



# Make alignment plot
plotMiro(paf.table = paf.subset, color.by = "direction", min.deletion.size = 50,
highlight.sv = "fill")

# Create annotation blocks for asip coding regions
# Create gene-like annotation
asip.gr <- GenomicRanges::GRanges(
```

```
            seqnames = 'chr2',
            ranges = IRanges::IRanges(start =
c(422954349,422955782,422958757),
                                  end = c(422954505,422955843,422958918)),
                                  ID = 'ASIP')


addAnnotation(ggplot.obj = plt, annot.gr = asip.gr, coordinate.space = "target",
           shape = 'rectangle', annotation.group = 'ID', fill.by = 'ID')


# Write to svg
ggsave("my_plot.svg")
```

**code S2: Script to plot eastern quoll mapping coverage across the ASIP locus**

```
# Script to plot eastern quoll mapping coverage
# across the ASIP locus

# Load in libraries
library(Gviz)
library(rtracklayer)

# Load in bedgraph files
coverage_data1 <- import("wildtype_1.bedgraph", format="bedGraph")
coverage_data2 <- import("wildtype_2.bedgraph", format="bedGraph")
coverage_data3 <- import("wildtype_3.bedgraph", format="bedGraph")
coverage_data4 <- import("melanistic_1", format="bedGraph")
coverage_data5 <- import("melanistic_2.bedgraph", format="bedGraph")
coverage_data6 <- import("melanistic_3.bedgraph", format="bedGraph")

# Load in asip annotation gff file
annotations <- import("asip.gff")

# Create datatracks for each sample
coverage_track1 <- DataTrack(coverage_data1, name="Wildtype 1",
col.histogram="#1F77B4", fill.histogram="#1F77B4", col.axis="black",
lwd.axis=1, drawAxis=TRUE)
coverage_track2 <- DataTrack(coverage_data2, name="Wildtype 2",
col.histogram="#1F77B4", fill.histogram="#1F77B4", col.axis="black",
lwd.axis=1, drawAxis=TRUE)
```

```
coverage_track3 <- DataTrack(coverage_data3, name="Wildtype 3",
col.histogram="#1F77B4", fill.histogram="#1F77B4", col.axis="black",
lwd.axis=1, drawAxis=TRUE)
coverage_track4 <- DataTrack(coverage_data4, name="Melanistic 4",
col.histogram="#D62728", fill.histogram="#D62728", col.axis="black",
lwd.axis=1, drawAxis=TRUE)
coverage_track5 <- DataTrack(coverage_data5, name="Melanistic 5",
col.histogram="#D62728", fill.histogram="#D62728", col.axis="black",
lwd.axis=1, drawAxis=TRUE)
coverage_track6 <- DataTrack(coverage_data6, name="Melanistic 6",
col.histogram="#D62728", fill.histogram="#D62728", col.axis="black",
lwd.axis=1, drawAxis=TRUE)


# Create a generegiontrack for asip
annotation_track <- GeneRegionTrack(annotations, name="Genes",
chromosome="chr2")

# Create a genomeaxistrack
genome_axis_track <- GenomeAxisTrack()

# Create an svg file for the plot
svg("coverage_plot.svg", width=10, height=7)

# Make the plot
plotTracks(list(annotation_track, genome_axis_track, coverage_track1,
coverage_track2,
           coverage_track3, coverage_track4, coverage_track5,
coverage_track6),
```

```
    from=422949000, to=422965000,
    type=c("hist", "g"),
    windowSize=5000,
    window=-1,
   lwd.axis=1,
    main="Multiple Coverage Tracks with Annotations (chr2:422949000-
422965000)")

# Write to svg
dev.off()
```

**code S3: Script to plot amino acid alignment of agreodont MC1R orthologs**

```
# Script to plot amino acid alignment of
# agreodont MC1R orthologs

# Load ggmsa
library(ggmsa)

# Load in alignment
protein_sequences <- "mc1r_cds.fna"

# Make plot
plot <- ggmsa(protein_sequences, start = 141, end = 180, char_width = 0.5,
seq_name = TRUE) + geom_seqlogo()

# Save plot to svg
ggsave("alignment_plot.svg", plot, width = 12, height = 6, dpi = 120000)
```
